# SIRT2 and lysine fatty acylation regulate the oncogenic activity of K-Ras4a

**DOI:** 10.1101/203638

**Authors:** Hui Jing, Xiaoyu Zhang, Stephanie A. Wisner, Xiao Chen, Nicole A. Spiegelman, Maurine E. Linder, Hening Lin

**Affiliations:** Department of Chemistry and Chemical Biology, Cornell University, Ithaca, NY 14853, USA; Department of Molecular Medicine, Cornell University College of Veterinary Medicine, Ithaca, New York 14853, USA; Howard Hughes Medical Institute, Department of Chemistry and Chemical Biology, Cornell University, Ithaca, NY 14853, USA

## Abstract

Ras proteins play vital roles in numerous biological processes and Ras mutations are found in many human tumors. Understanding how Ras proteins are regulated is important for elucidating cell signaling pathways and identifying new targets for treating human diseases. Here we report that one of the K-Ras splice variants, K-Ras4a, is subject to lysine fatty acylation, a previously under-studied protein post-translational modification. Sirtuin 2 (SIRT2), one of the mammalian nicotinamide adenine dinucleotide (NAD)-dependent lysine deacylases, catalyzes the removal of fatty acylation from K-Ras4a. We further demonstrate that SIRT2-mediated lysine defatty-acylation promotes endomembrane localization of K-Ras4a, enhances its interaction with A-Raf, and thus promotes cellular transformation. Our study identifies lysine fatty acylation as a previously unknown regulatory mechanism for the Ras family of GTPases that is distinct from cysteine fatty acylation. These findings highlight the biological significance of lysine fatty acylation and sirtuin-catalyzed protein lysine defatty-acylation.

## Introduction

Protein fatty acylation facilitates direct association of proteins with particular membranes in cells and plays a vital role in protein trafficking, cell signaling, protein-protein interactions, and protein activity^1–3^. Dysregulation of protein fatty acylation is implicated in human cancer and neurodegenerative diseases^3^. While early studies have focused on N-terminal glycine myristoylation and cysteine palmitoylation, little is known about lysine fatty acylation^2,3^. Although first reported over two decades ago, the biological function of protein lysine fatty acylation is not clear and to date, only a few proteins, such as tumor necrosis factor α (TNF-α) and interleukin 1-α, are known to be regulated by lysine fatty acylation^4–8^. The enzymes that catalyze the addition or removal of lysine fatty acylation were not known until recently when we and others found that several sirtuins, the nicotinamide adenine dinucleotide (NAD)-dependent protein lysine deacylase, could act as lysine defatty-acylase. We have previously reported that TNF-α ^9,10^ and exosome ^11^ secretion are regulated by sirtuin 6 (SIRT6)-catalyzed removal of lysine fatty acylation, demonstrating that lysine fatty acylation is reversible and physiologically important. Other sirtuin family proteins, SIRT1-3^12–16^ and SIRT7{Tong, 2017 #1557}, have also been found to efficiently remove fatty acyl groups from lysine residues *in vitro*, suggesting that lysine fatty acylation may be more prevalent. Therefore, we sought to identify other proteins that may be regulated by lysine fatty acylation.

Ras proteins are small GTPases that play important roles in numerous tumor-driving processes, including proliferation, differentiation, survival, cell cycle entry and cytoskeletal dynamics ^17^. They act as binary switches: they are active when GTP- bound, turning on specific signaling pathways by recruiting effector proteins, and inactive in the GDP- bound state ^17,18^. Guanine nucleotide exchange factors (GEFs) activate Ras by promoting GDP-GTP exchange, whereas GTPase-activating proteins (GAPs) inactivate Ras by promoting intrinsic GTP hydrolysis^17^. In mammals, *HRAS*, *NRAS*, and *KRAS* proto-oncogenes encode four proteins: H-Ras, N-Ras, K-Ras4a, and K-Ras4b. K-Ras4a and K-Ras4b are the two splice variants encoded by the *KRAS* gene. K-Ras4b has attracted most of the attention because it was assumed to be the more abundant and thus the more important K-Ras isoform mutated in human cancers. However, recent studies have revealed that K-Ras4a is widely expressed in many cancer cell lines and its level is similar to that of K-Ras4b in human colorectal tumors ^19,20^. A requirement for oncogenic K-Ras4a in lung carcinogenesis has also been demonstrated in mice ^21^. Thus, there is increasing interest in evaluating K-Ras4a as a therapeutic target and in investigating the regulation of K-Ras4a.

Ras proteins exert their functions at cellular membranes, where they interact with distinct effectors and activate downstream signaling ^18^. Ras proteins typically have two membrane-targeting signals at the C-terminal hypervariable regions (HVRs). All four Ras proteins are modified by cysteine farnesylation on their CaaX motif. H-Ras and N-Ras contain cysteine palmitoylation as the second membrane targeting signal, whereas K-Ras4b uses a polybasic region (PBR) (Fig. 1a). K-Ras4a possesses a hybrid membrane targeting motif: multiple lysine residues at the C-terminus (similar to K-Ras4b) as well as cysteine palmitoylation (Fig. 1a)^19,20^. As we set out to identify lysine fatty acylated proteins, the presence of multiple lysine residues in the Ras HVRs caught our attention. If the lysine residues function simply to promote membrane binding by electrostatics, why are there almost invariably lysine but not arginine residues in the HVRs? The prevalence of lysines in Ras HVRs suggests the possiblity that lysine residues are post-translationally modified by fatty acids. Thus, in this study, we set out to investigate whether Ras proteins are regulated by reversible lysine fatty acylation.

## Results

### H-Ras and K-Ras4a contain lysine fatty acylation

To examine whether the lysine residues in the Ras proteins could be fatty acylated, an alkyne-tagged fatty acid analog, Alk14, was used to metabolically label Ras proteins^9^. As shown in Fig. 1b, HEK293T cells transiently expressing FLAG-tagged H-Ras, N-Ras, K-Ras4a, or K-Ras4b were treated with Alk14 (50 μM). We ensured that the overexpression levels of different Ras proteins were similar (Fig. S1a).FLAG-tagged Ras proteins were immunoprecipitated and conjugated to rhodamine-azide (Rh-N_3_) using click chemistry to allow visualization of fatty acylation by in-gel fluorescence. Hydroxylamine (NH_2_OH) was then used to remove cysteine palmitoylation. Ras-related protein Ral-A (RalA)^22^ and Syntaxin-6 (STX6)^23^ were included as controls for the efficiency of NH_2_OH in removing cysteine palmitoylation. Quantification of the fluorescent signal revealed that NH_2_OH treatment removed over 95% of the fatty acylation from RalA or STX6. However, H-Ras, N-Ras and K-Ras4a retained 20%, 13% and 47% relative NH_2_OH-resistant fatty acylation over total fatty acylation, respectively, whereas K-Ras4b did not show Alk14 labeling either before or after NH_2_OH treatment (Fig. 1c). These data suggest that H-Ras, N-Ras, and K-Ras4a might possess non-cysteine fatty acylation.

To determine whether the NH_2_OH-resistant fatty acylation could be attributed to the lysine residues in the HVRs of H-Ras, N-Ras, and K-Ras4a, we mutated these lysine (K) residues to arginine (R) and examined the fatty acylation of the WT and KR mutants. The H-Ras 3KR (K167/170/185R) mutant (Fig. 1d), N-Ras 2KR (K169/170) mutant (Fig. S1b), and the K-Ras4a 3KR (K182/184/185R) mutant but not K-Ras4a 4KR (K169/170/173/176) mutant (Fig. 1e & f) all displayed greatly reduced NH_2_OH-resistant fatty acylation, implying that H-Ras, N-Ras and K-Ras might be fatty acylated on the lysine residues in their HVRs. Moreover, K-Ras4a-7KR (4KR & 3KR) showed comparable NH_2_OH-resistant fatty acylation level to the 3KR mutant, suggesting that K182/184/185 of K-Ras4a might be fatty acylated. We then utilized mass spectrometry (MS) to directly identify the lysine fatty acylation of FLAG-H-Ras, -N-Ras or -K-Ras4a extracted from Alk14-treated HEK293T cells with tryptic digestion. This allowed us to identify H-Ras K170 (Fig. 1g) and K-Ras4a K182 (Fig. 1h) as being modified, confirming lysine fatty acylation of H-Ras and K-Ras4a. Our attempt to identify N-Ras lysine fatty acylation by MS was not successful possibly because the tryptic peptide with lysine fatty acylation was less abundant (Fig. 1c), too short (MK_acyl_K) or too hydrophobic (K_acyl_LNSSDDGTQGC_cam_MGLPC_prenyl,ome_)^1,2,24^.

**Figure 1.**
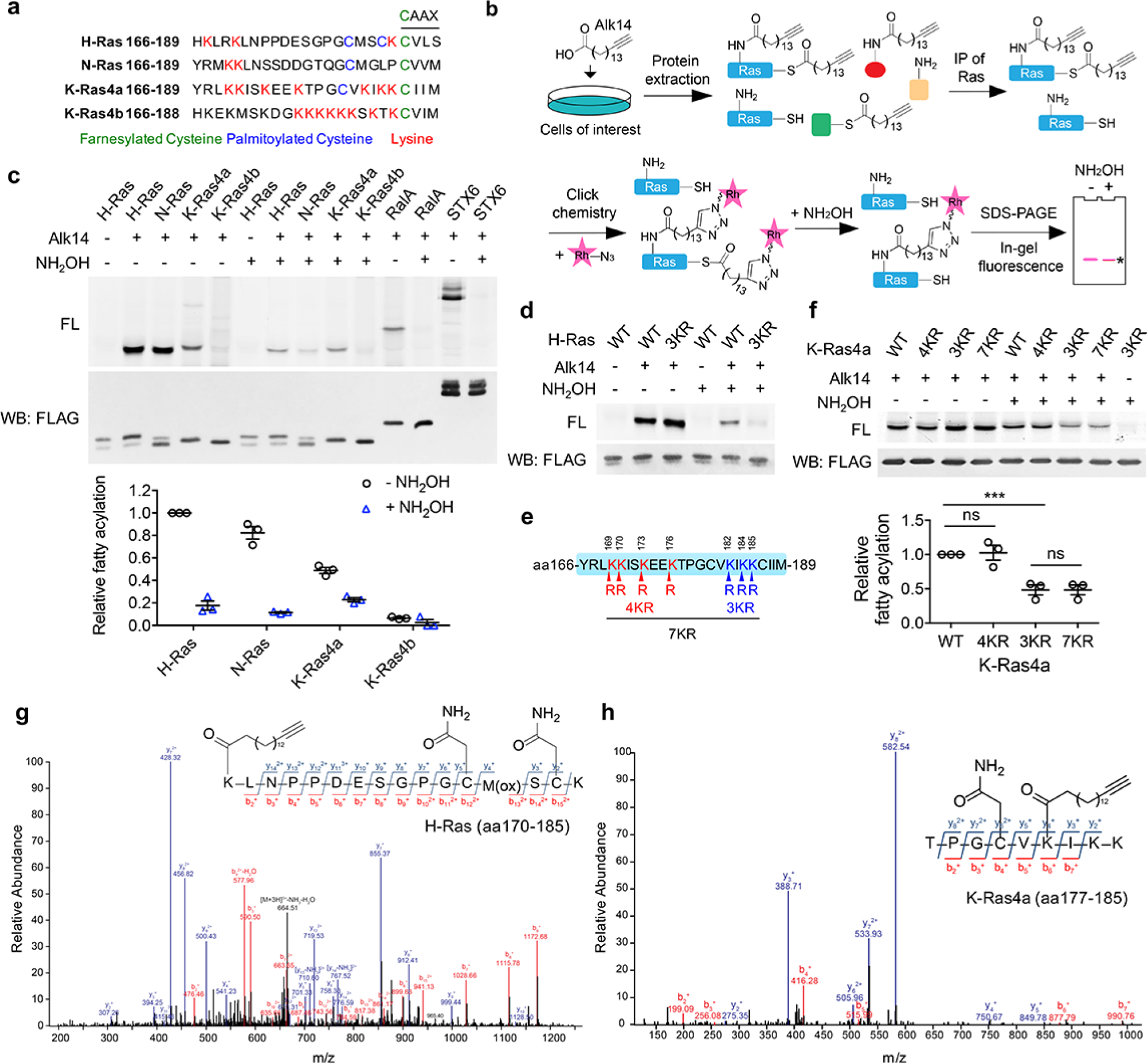
H-Ras and K-Ras4a contain lysine fatty acylation. (**a**) Amino acid sequences of the HVRs of Ras proteins. (**b**) Scheme showing the Alk14 metabolic labeling method to study lysine fatty acylation. (**c**) In-gel fluorescence detection of the fatty acylation levels of Ras proteins, RalA and STX6 in HEK293T cells (top panel), and quantification of the relative fatty acylation levels (bottom panel). The fatty acylation level of H-Ras without NH_2_OH treatment was set to 1. (**d**) In-gel fluorescence showing the fatty acylation levels of H-Ras WT and 3KR mutant without or with NH_2_OH treatment. (**e**) Scheme showing the lysine to arginine mutants (4KR, 3KR, and 7KR) used to identify potential fatty acylation sites. (**f**) In-gel fluorescence showing the fatty acylation levels of K-Ras4a WT and KR mutants without or with NH2OH treatment (top panel) and quantification of fatty acylation levels with NH2OH treatment relative to that of K-Ras4a WT (bottom panel). (**g,h**) Tandem mass (MS/MS) spectrum of triply charged H-Ras (**g**) and K-Ras4a (**h**) peptides with Alk14-modification on K170 and K182, respectively. The b- and y-ions are shown along with the peptide sequence. The cysteine residues were carbamidomethylated due to iodoacetamide alkylation during sample preparation and methionine was oxidized. FL, fluorescence; WB, western blot. Statistical evaluation was by unpaired two-tailed Student’s t test. Error bars represent SEM in three biological replicates. \*\*\**P* < 0.001; ns, not significant. Representative images from three independent experiments are shown.

### SIRT2 catalyzes the removal of lysine fatty acylation from K-Ras4a

Several mammalian sirtuins, including SIRT1, SIRT2, SIRT3, and SIRT6, can efficiently remove fatty acyl groups from protein lysine residues *in vitro* (SIRT1, SIRT2, and SIRT3)^25^ or *in vivo* (SIRT6)^9,12–14,25^. So we next investigated whether any of the these sirtuins could remove lysine fatty acylation from H-Ras or K-Ras4a and therefore regulate their function. We incubated H-Ras or K-Ras4a isolated from Alk14-treated HEK293T cells with purified recombinant sirtuins without or with NAD *in vitro* and examined the H-Ras or K-Ras4a fatty acylation level by in-gel fluorescence after click chemistry. Incubation of H-Ras or K-Ras4a with *Plasmodium falciparum* Sir2A (PfSir2A), a sirtuin family member with robust lysine defatty-acylase activity^26^, resulted in the removal of most of the NH_2_OH-resistant fatty acylation from H-Ras and K-Ras4a in the presence of NAD (Fig. S2a). This result further confirmed that the NH_2_OH-resistant fatty acylation is mainly from lysine residues and indicated that lysine fatty acylation of H-Ras and K-Ras4a is reversible. Furthermore, SIRT2, but not SIRT1, 3, or 6, slightly decreased the lysine fatty acylation signal of H-Ras (Fig. 2a); SIRT1 and SIRT2, but not SIRT3 and SIRT6, removed lysine fatty acylation from K-Ras4a. Notably, SIRT2 showed better activity than SIRT1 on K-Ras4a lysine fatty acylation (Fig. 2b). In contrast, SIRT1 and SIRT2 showed little effect on the fatty acylation of K-Ras4a-3KR (Fig. S2b), which exhibited significantly lower lysine fatty acylation than K-Ras4a-WT (Fig. 1f), suggesting that SIRT1 and SIRT2 do not possess cysteine defatty-acylase activity. An HPLC-based *in vitro* activity assay also revealed that SIRT2 was unable to remove the cysteine myristoyl group from a K-Ras4a-C180myr peptide (Fig. S2c). Furthermore, knockdown (KD) of SIRT2 in HEK293T cells did not affect lysine fatty acylation of H-Ras (Fig. 2c & f), whereas KD of SIRT2 but not SIRT1 significantly increased lysine fatty acylation of K-Ras4a compared with control (Ctrl) KD (Fig. 2d, e & f). We also noted that SIRT2 KD did not affect fatty acylation of N-Ras (Fig. S2d). Taken together, these results illustrate that K-Ras4a is a lysine defatty-acylation substrate for SIRT2 in cells, but H-Ras and N-Ras are not.

**Figure 2.**
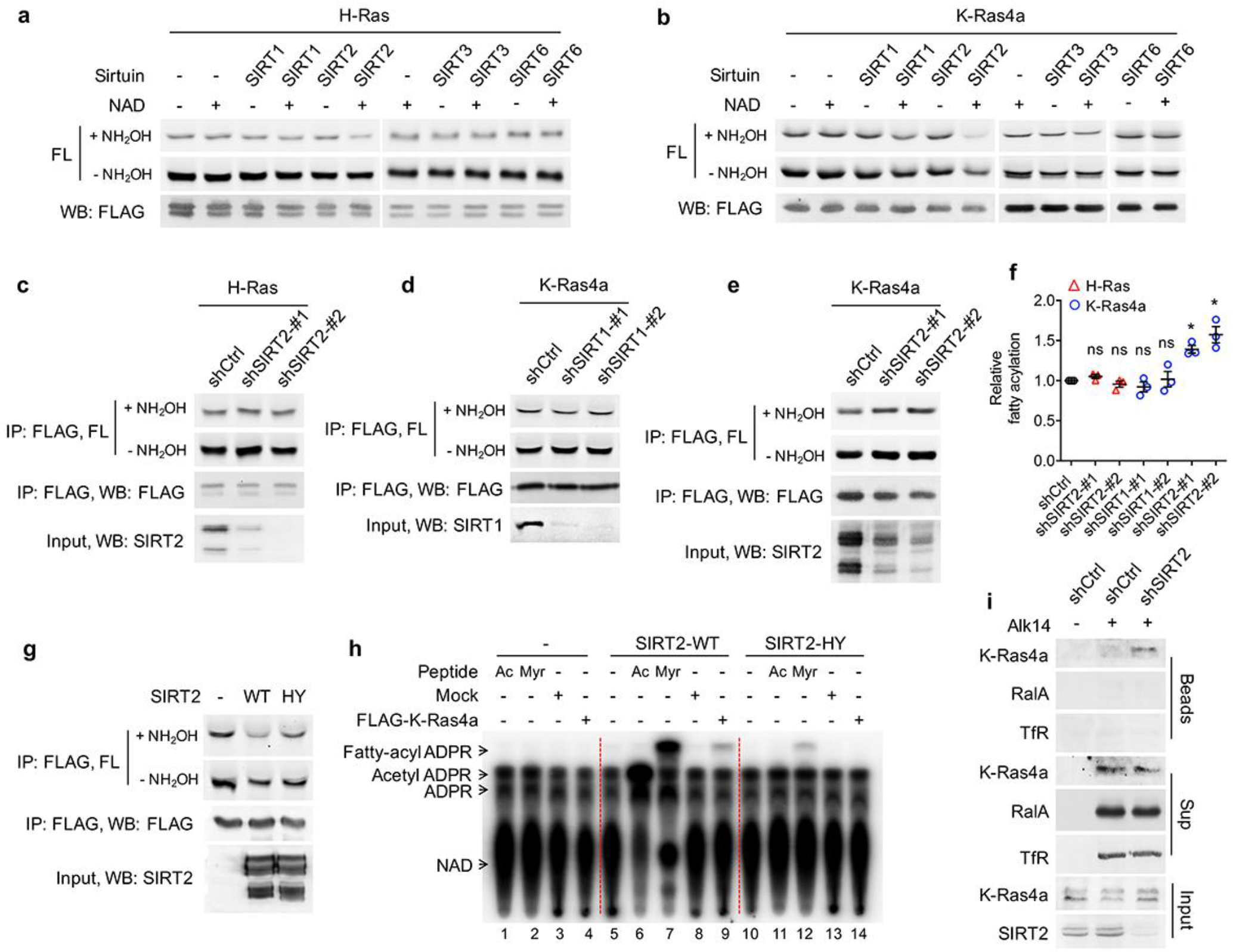
SIRT2 removes lysine fatty acylation from K-Ras4a. (**a, b**) In-gel fluorescence detection of fatty acylation of H-Ras (a) and K-Ras4a (b) treated with 5 μM of SIRT1, 2, 3 and 6 without or with 1 mM of NAD *in vitro.* (**c**) Effect of SIRT2 KD on the fatty acylation level of H-Ras in HEK293T cells. (**d, e**) Effect of SIRT1 KD (d) and SIRT2 KD (e) on the fatty acylation level of K-Ras4a in HEK293T cells. (**f**) Quantification of the fatty acylation levels with NH_2_OH treatment in (c), (d) and (e). The fatty acylation level in the corresponding Ctrl KD was set to 1. (**g**) Effect of overexpressing SIRT2-WT and SIRT2-HY catalytic mutant on K-Ras4a fatty acylation level. (**h**) Fatty acylated lysine in K-Ras4a detected by formation of ^32^P-labeled fatty acyl-ADPR using ^32^P-NAD. (**i**) Lysine fatty acylation of endogenous K-Ras4a in Ctrl and **SIRT2** KD (by shSIRT2-#2) HCT116 cells detected by Alk14 labeling and biotin pull-down. Sup, supernatant; Ac, acetyl-H3K9; Myr, myristoyl-H3K9. Statistical evaluation was by unpaired two-tailed Student's t test. Error bars represent SEM in three biological replicates. \**P* < 0.05; ns, not significant. Representative images from three independent experiments are shown.

We next further validated that K-Ras4a is regulated by SIRT2-mediated defatty-acylation. We utilized the SIRT2-H187Y (HY) mutant, which has previously been shown to be catalytically dead in lysine deacetylation^27^, as a negative control. An HPLC-based *in vitro* assay demonstrated that the H187Y mutation dramatically decreased SIRT2 defatty-acylation activity, while it completely abolished its deacetylation activity (Fig. S2e). Co-expression of SIRT2 with K-Ras4a in HEK293T cells substantially decreased K-Ras4a lysine fatty acylation, whereas co-expression of SIRT2-HY had much less effect (Fig. 2g), suggesting that K-Ras4a defatty-acylation requires SIRT2 catalytic activity. Interestingly, our finding that mutation of the catalytic histidine residue did not completely abolish sirtuin enzymatic activity is not without precedent. For example, mutating the catalytic histidine of bacterial Sir2Tm^28^, yeast HST2^29^, and human SIRT6^11^ also retained some catalytic activity.

To investigate whether K-Ras4a could also be regulated by SIRT2 through deacetylation, we examined its acetylation level using a pan-specific acetyl lysine antibody. Acetylation was not detected on K-Ras4a in either Ctrl KD or SIRT2 KD cells without or with histone deacetylases (HDAC) inhibitor Trichostatin A (TSA) (Fig. S2f). We also searched our K-Ras4a MS data and did not find any peptides with lysine acetylation, indicating that SIRT2 likely does not regulate K-Ras4a via deacetylation.

With SIRT2 as a tool, we further confirmed the existence of lysine fatty acylation on K-Ras4a in cells that were not treated with Alk14. We used a previously developed assay that relies on ^32^P-NAD to detect sirtuin-catalyzed deacylation reactions^30^. When histone H3K9 acetyl (Ac) and myristoyl (Myr) peptides were incubated with SIRT2-WT in the presence of ^32^P-NAD, the formation of the acyl-ADPR product could be detected by autoradiography after separation using thin-layer chromatography (TLC) (Fig. 2h, lanes 6 & 7). In contrast, the SIRT2-HY mutant only generated a tiny amount of the acyl-ADPR product (Fig. 2h, lanes 11 & 12). When K-Ras4a isolated from HEK293T cells was treated with SIRT2-WT in the presence of ^32^P-NAD, a spot corresponding to fatty acyl-ADPR but not acetyl-ADPR was detected (Fig. 2h, lane 9). Control reactions without SIRT2 (Fig. 2h, lane 4) or with the HY mutant (Fig. 2h, lane 14) did not generate the fatty acyl-ADPR product. These results demonstrate that K-Ras4a contains lysine fatty acylation that can be removed by SIRT2 in the absence of Alk14 supplementation. A peptide carrying palmitoylation, but not myristoylation, on K182 of FLAG-tagged K-Ras4a was detected by MS (Fig. S2g), demonstrating that palmitoylation is the major native lysine acylation of K-Ras4a.

We then investigated whether endogenous K-Ras4a is also regulated by SIRT2-catalyzed lysine defatty-acylation. For this purpose, we used the HCT116 human colorectal cancer cell line, in which K-Ras4a was shown to be expressed^19^. Since the commercial antibody against K-Ras4a did not immunoprecipitate K-Ras4a, we enriched fatty acylated proteins labeled with Alk14 as previously described^31^, and detected fatty acylated K-Ras4a using a K-Ras4a-specific antibody. HCT116 cells with Ctrl KD or SIRT2 KD were cultured in the presence of Alk14. Proteins were then extracted and a biotin affinity tag was attached to the Alk14-labeled proteins with click chemistry. The biotin-conjugated proteins were pulled down using streptavidin beads, and subsequently washed with 1% SDS to disrupt protein-protein interaction. Proteins that were only fatty acylated on cysteine residues were then released from the streptavidin beads into the supernatant (Sup) *via* NH_2_OH treatment, while proteins with lysine fatty acylation were retained. As shown in Fig. 2i, RalA (Fig. 1c) and transferrin receptor (TfR)^32^, which are predominantly cysteine fatty acylated, were present in the supernatant but barely detectable from the streptavidin beads, indicating that the NH_2_OH treatment was effective. In Ctrl KD cells, K-Ras4a was mainly detected in the supernatant. However, in the SIRT2 KD cells, K-Ras4a was detected both on the streptavidin beads and in the supernatant, indicating that endogenous K-Ras4a possesses lysine fatty acylation that is regulated by SIRT2.

By immunoprecipitation of total Ras protein from Alk14-treated HCT116 cells using a pan-Ras (Y13-259) antibody, we found that endogenous Ras proteins exhibited NH_2_OH-resistant fatty acylation (Fig. S3a). Moreover, SIRT2 KD increased the NH_2_OH-resistant fatty acylation of Ras proteins (Fig. S3b). Since SIRT2 KD did not affect lysine fatty acylation of overexpressed H-Ras (Fig. 2c & f) and N-Ras (Fig. S2d), the data suggested that the increase in total Ras lysine fatty acylation observed in SIRT2 KD cells can be attributed to K-Ras4a lysine fatty acylation. This result, together with the detection of K-Ras4a lysine fatty acylation by Alk14 biotinylation, further supports that endogenous K-Ras4a is lysine fatty acylated and is regulated by SIRT2-mediated lysine defatty-acylation. We also performed MS analysis of endogenous Ras immunoprecipitated from HCT116 cells treated with SIRT2 shRNA and Alk14. We identified a peptide with a primary mass matching the Alk14-modified K-Ras4a aa177-185 peptide, whose exact m/z and isotope pattern were the same as those of overexpressed K-Ras4a (Fig. S3c). However, this primary mass did not trigger MS2, which was likely due to low peptide abundance (Fig. S3c & d). It has been shown that K-Ras has a much lower expression level than H-Ras and N-Ras because of its rare codon bias^33,34^.

SIRT2 was reported to reside predominately in the cytoplasm^35,36^. The regulation of K-Ras4a lysine fatty acylation by SIRT2 suggested that SIRT2 might also exist at cellular membranes, where K-Ras4a mainly resides. Indeed, by subcellular fractionation, we found that SIRT2 was present in both soluble and membrane fractions (Fig. S2h). Co-immunoprecipitation (co-IP) revealed K-Ras4a associated with endogenous SIRT2 (Fig. S2i). These results further support that K-Ras4a is a lysine defatty-acylase substrate for SIRT2.

### Mapping the fatty acylated lysine residues regulated by SIRT2

MS results suggested that K182 was preferentially fatty acylated lysine on K-Ras4a. However, the lysine 182 to arginine mutant (K182R) exhibited similar lysine fatty acylation levels to that of WT (Fig. 3a). As the 3KR (K182/184/185R) but not the 4KR (K169/170/173/176R) mutant significantly decreased K-Ras4a lysine fatty acylation (Fig. 1f), we also mutated K184 and K185 to arginine individually. Neither the K184R nor K185R mutation decreased lysine fatty acylation as the 3KR mutant did (Fig. 3a). These results suggested that K182, 184 and 185 were likely to be modified redundantly. We suspected that it was hard to pinpoint the exact modification site by mutagenesis because the K182R mutation might enhance fatty acylation on the other two nearby lysine residues. To test this hypothesis, we performed MS analysis of FLAG-K-Ras4a-K182A extracted from Alk14-treated HEK293T cells. We tested the K182A instead of K182R mutant because the K182R mutant would produce a tryptic peptide that is too short to be detected. As expected, the K182A mutation did not affect overall level of K-Ras4a lysine fatty acylation (Fig. 3b). A peptide (amino acids 177-185) fatty acylated on K184 was detected by MS (Fig. 3b), which agrees with our hypothesis. It was likely that K185 could also be fatty acylated for the K182R mutant, because the K182/185R mutant slightly but significantly decreased lysine fatty acylation levels compared with the K182R and K185R single mutants (Fig. 3a). The K185 fatty acylation was not detected by MS most likely because the modified tryptic peptide was too short and hydrophobic (K_fatty-acyl_C_prenyl,oMe_). Overall, these data indicate that K182/184/185 are fatty acylated redundantly and that the 3KR mutation is needed to abolish lysine fatty acylation on the C-terminus of K-Ras4a.

**Figure 3.**
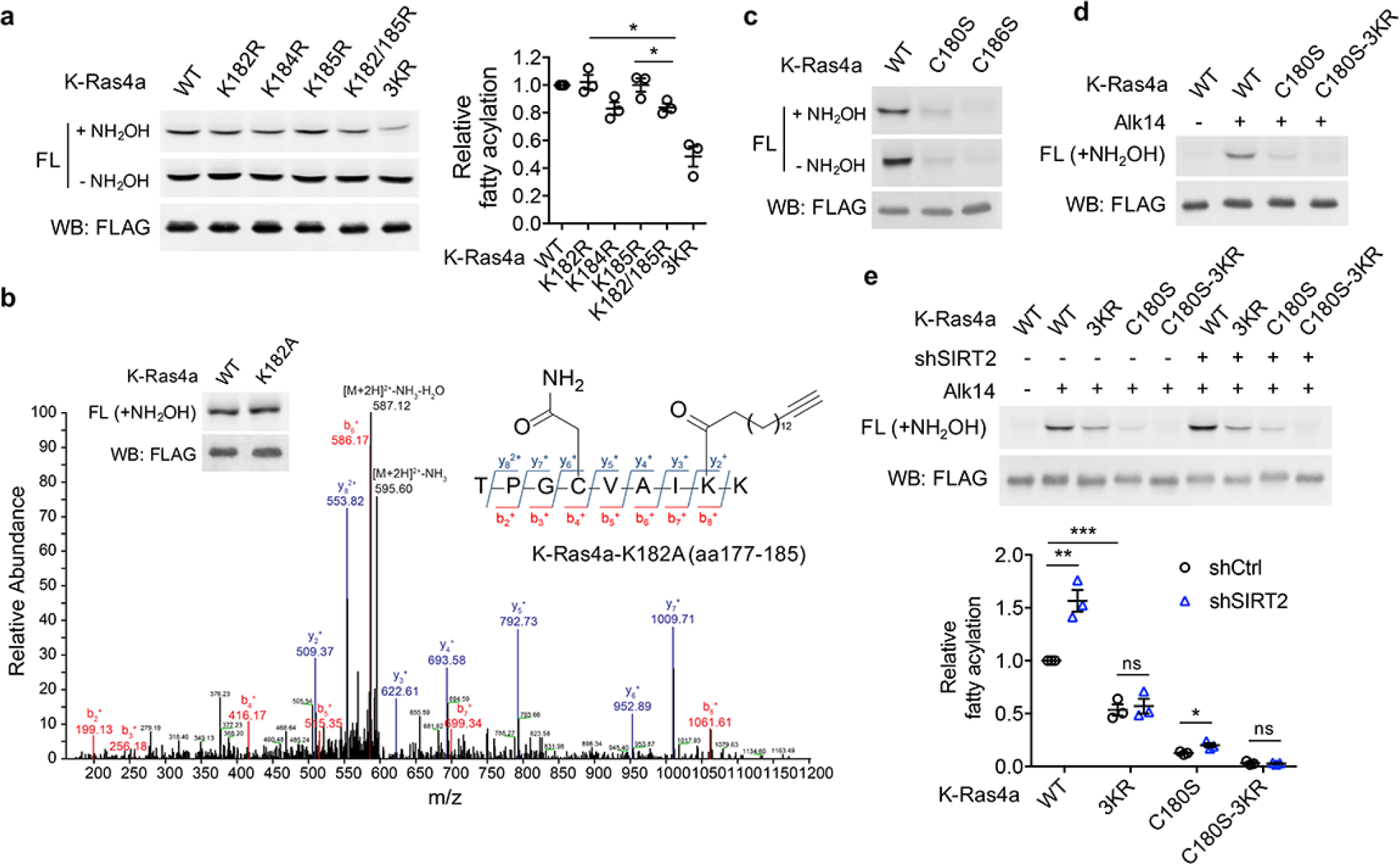
SIRT2 regulates lysine fatty acylation of K-Ras4a on K182/184/185. (**a**) Fatty acylation levels of K-Ras4a WT, K182R, K184R, K185R, K182/185R, and 3KR by in-gel fluorescence (left panel) and quantification of fatty acylation levels after NH_2_OH treatment relative to that of K-Ras4a WT (right panel). (**b**) MS/MS spectrum of triply charged K-Ras4a-K182A peptide with Alk14 modification on K184. The b- and y-ions are shown along with the peptide sequence. The cysteine residue was carbamidomethylated due to iodoacetamide alkylation during sample preparation. Fatty acylation levels of K-Ras4a WT and K182A with NH_2_OH were also shown. (**c**) Fatty acylation levels of K-Ras4a WT, C180S, and C186S. (**d**) Fatty acylation levels of K-Ras4a WT, C180S and C180S-3KR after NH_2_OH treatment. (**e**) Fatty acylation levels of K-Ras4a WT, 3KR, C180S and C180S-3KR after NH_2_OH treatment in Ctrl and SIRT2 KD (by shSIRT2-#2) HEK293T cells. Quantification of the fluorescent intensity relative to K-Ras4a WT is shown in the bottom panel. Statistical evaluation was by unpaired two-tailed Student’s t test. Error bars represent SEM in three biological replicates. \**P* < 0.05; \*\**P* < 0.01; \*\*\**P* < 0.001. Representative images from three independent experiments are shown.

K-Ras4a has been shown to be prenylated on cysteine 186 and palmitoylated on cysteine 180^19^. To examine whether cysteine prenylation or palmitoylation play a role in lysine fatty acylation, we generated cysteine-to-serine C180S and C186S mutants. Mutation of the prenylcysteine (C186S) completely abolished the fatty acylation of K-Ras4a (Fig. 3c), which is consistent with the model that prenylation of the cysteine on the CaaX motif of Ras proteins is required for the subsequent fatty acylation^37^. On the other hand, mutation of the palmitoylated cysteine (C180S) led to a substantial, but not a complete, loss of K-Ras4a lysine fatty acylation (Fig. 3c). The fatty acylation on the C180S mutant was NH_2_OH-resistant and was abolished by combining the C180S and 3KR mutations (Fig. 3d), implying that the C180S mutant was fatty acylated on K182/184/185. These data suggest that cysteine palmitoylation might play an important but nonessential role in the occurrence of lysine fatty acylation. It is possible that cysteine palmitoylation facilitates the lysine fatty acyl transfer reaction, or the delivery of K-Ras4a to where lysine fatty acylation occurs.

We next assessed whether SIRT2 regulates fatty acylation of K-Ras4a on K182/184/185. SIRT2 removed lysine fatty acylation from K-Ras4a WT, the 4KR mutant and the C180S mutant, but not the 3KR mutant *in vitro* (Fig. S4a & b). SIRT2 KD in HEK293T cells increased lysine fatty acylation of K-Ras4a WT and the C180S mutant, but not the 3KR and C180S-3KR mutants (Fig. 3e), indicating that fatty acylation on K182/184/185 is regulated by SIRT2.

### Lysine fatty acylation regulates subcellular localization of K-Ras4a

We next set out to study the effect of lysine fatty acylation on K-Ras4a. A variety of PTMs on Ras proteins, such as cysteine palmitoylation ^38,39^, phosphorylation^40,41^ and ubiquitination^42^, function to deliver the molecule to the right place within the cell. We hypothesized that lysine fatty acylation may also be critical for the correct subcellular distribution of K-Ras4a. To test this hypothesis, we fused *Aequorea coerulescens* Green Fluorescent Protein (GFP) to the N-terminus of K-Ras4a WT and the 3KR mutant and performed live imaging with confocal microscopy in Ctrl and SIRT2 KD HEK293T cells to visualize K-Ras4a localization. The levels of over-expressed K-Ras4a WT and 3KR were equal in Ctrl and SIRT2 KD cells (Fig. S5a). We also imaged cells with similar GFP intensity under the same settings to avoid potential false positive observations caused by different levels of expression. In Ctrl KD cells, both K-Ras4a WT and the 3KR mutant displayed predominant localization to the plasma membrane (PM). However, the presence of 3KR on intracellular puncta was noticeably more pronounced compared to WT. SIRT2 KD decreased the intracellular punctate-localized K-Ras4a WT compared to Ctrl KD, whereas it had no effect on the punctate localization of the 3KR mutant (Fig. 4a & b), indicating that the effect of the 3KR mutation on K-Ras4a localization was due to lack of lysine fatty acylation. Similar effects of the 3KR mutation were obtained for K-Ras4a in HCT116 cells and for oncogenic K-Ras4a-G12V, which exhibited a comparable lysine fatty acylation level to K-Ras4a WT (Fig. S5b) in NIH3T3 cells (Fig. S5c & d). On the other hand, SIRT2 KD did not affect the intracellular punctate localization of H-Ras (Fig. 4a & b), which is consistent with our observation that H-Ras was not regulated by SIRT2 through lysine defatty-acylation. Taken together, these data indicate that lysine fatty acylation inhibits the intracellular punctate localization of K-Ras4a and SIRT2 promotes this localization by defatty-acylation. In addition, the K-Ras4a-C180S mutant that lacks cysteine palmitoylation and the majority of lysine fatty acylation extensively localized to internal membranes, which was distinct from the punctate localization of the 3KR mutant that is deficient in lysine fatty acylation but retains cysteine palmitoylation (Fig. S5e & f, Movie S1, 2, 3). This implies that cysteine palmitoylation might facilitate the punctate localization of K-Ras4a in the absence of lysine fatty acylation, while lysine fatty acylation inhibits it.

It has been shown that Ras proteins associate with and signal from endomembrane compartments, including the endoplasmic reticulum (ER), Golgi, endosomes and lysosome ^18,42–48^. Therefore, we next set out to identify the endomembrane compartments where lysine defatty-acylated K-Ras4a is localized. We performed colocalization analyses with a series of membrane compartment markers. Compared with K-Ras4a WT, the 3KR mutant exhibited more pronounced cytoplasmic colocalization with *trans*-Golgi network (TGN) marker STX6, early endosome marker EEA1, recycling endosome marker Rab11, and lysosome marker LAMP1 (Fig. 4c & d), but not with the ER marker Sec61, *trans*-Golgi marker GalT and late endosome marker Rab7 (Fig. S5g & h). Time-lapse confocal imaging also revealed that K-Ras4a-3KR displayed more internalization from the plasma membrane into punctate structures than did the WT (Supplemental Movie S1 & 2). These results suggest that removal of lysine fatty acylation from K-Ras4a promotes its localization to endomembranes in endocytic pathways, by which it may be routed from early endosome to the lysosome for degradation and to the TGN or recycling endosomes to return to the plasma membrane ^49^.

**Figure 4.**
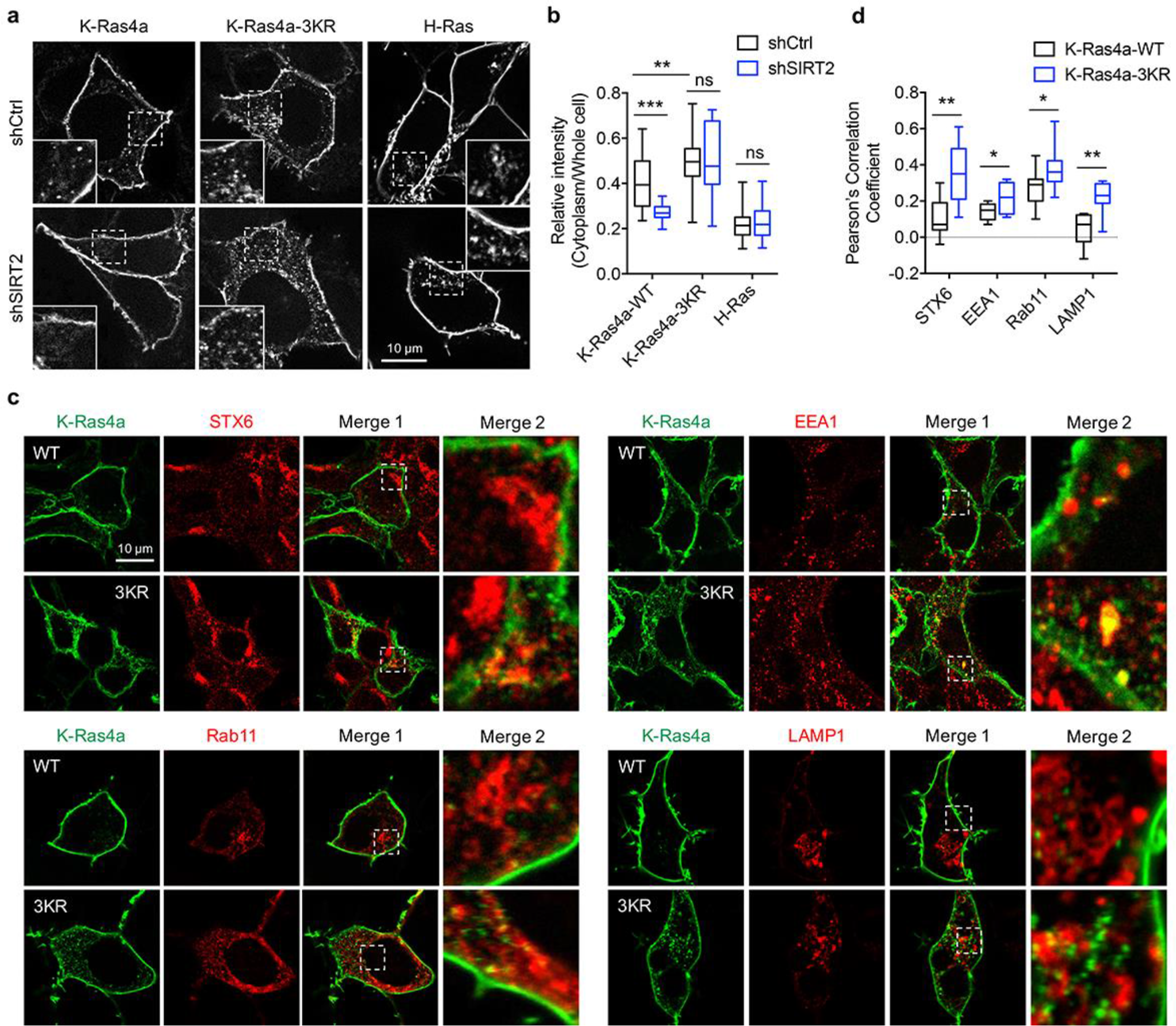
Lysine fatty acylation regulates subcellular localization of K-Ras4a. (**a**) Confocal images showing subcellular localization of GFP-K-Ras4a WT, 3KR, and GFP-H-Ras in HEK293T cells with Ctrl or SIRT2 KD (by shSIRT2-#2). Insets are magnifications of the regions enclosed by the white dashed squares. (**b**) Statistical analyses of the relative cytoplasm to whole cell intensity of K-Ras4a WT, 3KR, and H-Ras from (a) (n = 16, 16, 16, 16, 21, 21 for each sample from left to right, respectively). (**c**) Images showing the colocalization of GFP-K-Ras4a WT or 3KR with STX6, EEA1, Rab11, and LAMP1 in HEK293T cells. Merge 2 shows the magnified white dashed squares-enclosed regions in Merge 1. (**d**) Statistical analyses of the cytoplasmic colocalization of K-Ras4a or -3KR with the indicated intracellular membrane markers from (c) using Pearson's coefficient (n = 11, 11, 11, 11, 17, 17, 10, 10 cells for each sample from left to right, respectively). Statistical evaluation was by two-way ANOVA. *Centre line* of the box plot represents the mean value, box represents the 95 % confidence interval, and whiskers represent the range of the values. \**P* < 0.05; \*\**P* < 0.01; \*\*\**P* < 0.001; ns, not significant. Representative images are shown.

### Lysine fatty acylation regulates transforming activity of K-Ras4a

We next investigated whether lysine fatty acylation also affects the function of K-Ras4a. We assessed the ability of constitutively active K-Ras4a-G12V and the K-Ras4a-G12V-3KR mutant to enable anchorage-independent growth, promote proliferation in monolayer cultures and stimulate migration in Ctrl and Sirt2 KD cells. In Ctrl KD cells, expression of K-Ras4a-G12V-3KR resulted in significantly more colony formation on soft agar than did expression of K-Ras4a-G12V. Furthermore, Sirt2 KD caused a greater decrease in colony formation induced by K-Ras4a-G12V (75% decrease) than by K-Ras4a-G12V-3KR (45% decrease) (Fig. 5). Additionally, Sirt2 KD more potently inhibited K-Ras4a-G12V-mediated colony formation than H-Ras4a-G12V-mediated colony formation (Fig. 5), consistent with the fact that SIRT2 regulates lysine fatty acylation of K-Ras4a but not H-Ras or K-Ras4a 3KR. Thus, lysine fatty acylation inhibits the ability of K-Ras4a-G12V to induce anchorage-independent growth of cells and SIRT2 promotes it through defatty-acylation. One caveat of the result, however, was that Sirt2 KD still decreased the colony formation induced by K-Ras4a-G12V-3KR or H-Ras-G12V, whose lysine fatty acylation was not regulated by SIRT2. This is not unexpected because SIRT2 is known to exert tumor-promoting functions by deacetylating various targets^50–58^. Thus, the effect of Sirt2 KD on K-Ras4a-G12V-3KR- and H-Ras-induced transformation might be attributed to other substrates for SIRT2.

In monolayer cultures, NIH3T3 cells expressing K-Ras4a-G12V-3KR displayed a higher proliferation rate than those expressing K-Ras4a-G12V. Sirt2 KD inhibited the proliferation of the NIH3T3-K-Ras4a-G12V cells (47% inhibition) slightly more than that of the NIH3T3-K-Ras4a-G12V-342 3KR (34% inhibition) cells (Fig. S6a). Thus, lysine fatty acylation negatively regulates K-Ras4a-G12V-induced cell proliferation under monolayer culture conditions, but the effect was smaller than that on anchorage-independent growth (Fig. 5). Results from transwell migration assays revealed that the 3KR mutation did not affect the capability of K-Ras4a-G12V to induce cell migration. Consistent with this finding, Sirt2 KD decreased K-Ras4a-G12V and K-Ras4a-G12V-3KR-mediated cell migration similarly (Fig. S6b & c). Therefore, lysine fatty acylation does not affect the ability of K-Ras4a-G12V to stimulate cell migration.

**Figure 5.**
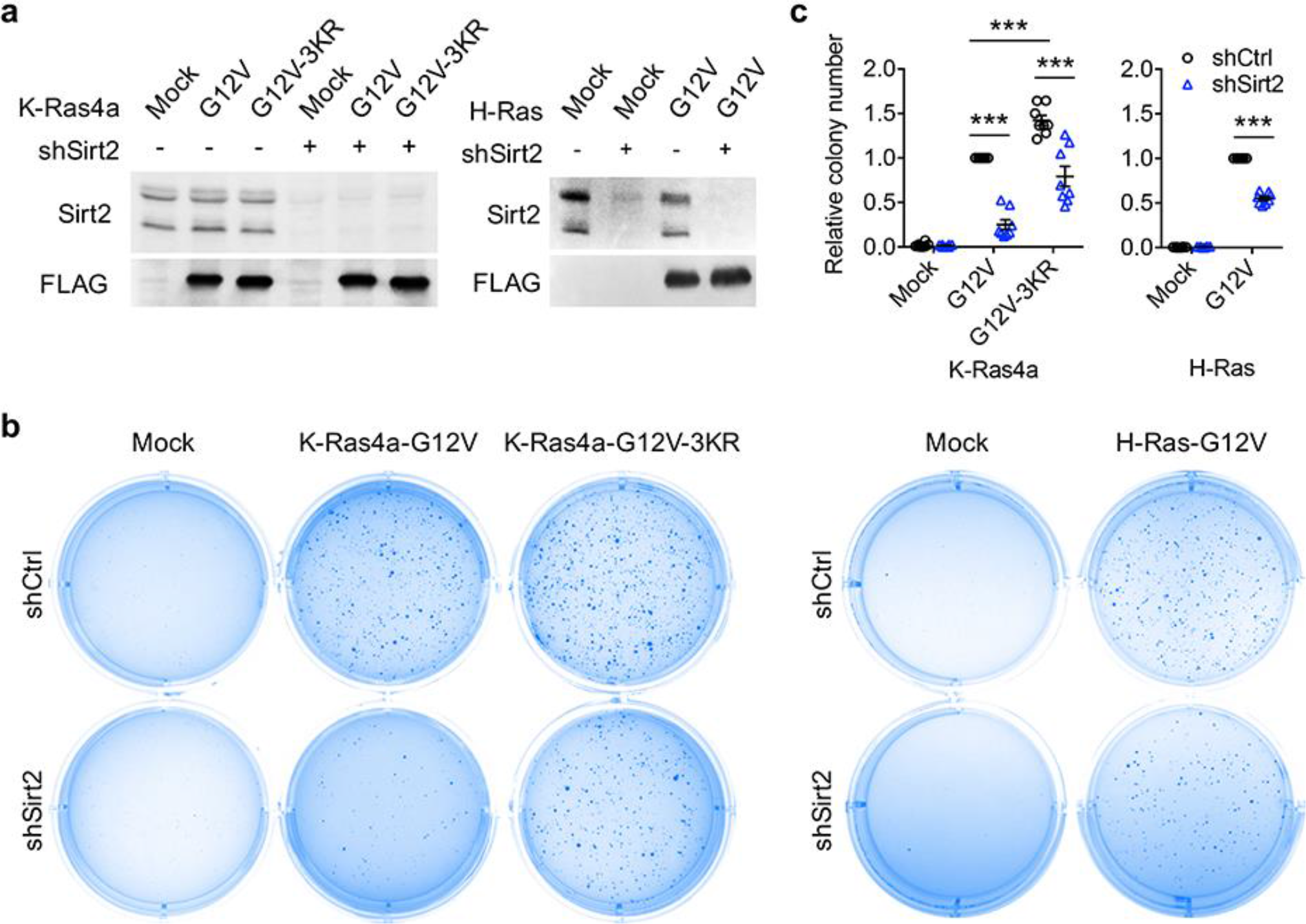
SIRT2-dependent lysine defatty-acylation increases K-Ras4a transforming activity. (**a**) Representative western blot analyses of Sirt2, FLAG-K-Ras4a-G12V, FLAG-K-Ras4a-G12V-3KR, FLAG-H-Ras-G12V protein levels in NIH3T3 cells with Ctrl or Sirt2 KD used in (**b** and **c**). (**b**) Anchorage-independent growth of NIH 3T3 cells stably expressing Mock, K-Ras4a-G12V, -G12V-3KR or H-Ras-G12V with Ctrl or Sirt2 KD. (**c**) Quantification of the colony numbers in (**b**) relative to that of the cells expressing K-Ras4a-G12V-shCtrl or H-Ras-G12V-shCtrl. Statistical evaluation was by unpaired two-tailed Student’s t test. Error bars represent SEM in eight biological replicates or as indicated. \*\*\**P* < 0.001. Representative images (a, b) from at least three independent experiments are shown.

### A-Raf is involved in the regulation of K-Ras4a by lysine fatty acylation

The dynamic regulation of Ras localization is known to be closely coupled to its signaling output ^43,59,60^. We decided to further explore the molecular mechanism underlying the regulation of K-Ras4a-mediated transformation by lysine fatty acylation. We first sought to examine whether lysine fatty acylation affects K-Ras4a activation by a pull-down assay with the Ras-binding domain (RBD) of Raf1, which only binds to the GTP-bound form of Ras^43^. Neither the 3KR mutation nor SIRT2 KD affected EGF-stimulated GTP loading of K-Ras4a (Fig. S7a & b) or the constitutively GTP-loaded state of K-Ras4a-G12V (Fig. S7c & d). We then determined whether K-Ras4a at endomembranes exists in its GTP-bound state using DsRed-RBD (DsRed fused to the N-terminus of RBD) as a probe. Notably, we observed more colocalization of DsRed-RBD with K-Ras4a on intracellular puncta in cells expressing the 3KR mutant (Fig. S7e, f, g) than in cells expressing K-Ras4a WT. Furthermore, SIRT2 KD decreased the colocalization of DsRed-RBD with K-Ras4a WT at intracellular puncta, but not with the 3KR mutant (Fig. S7e).

The results above suggest that SIRT2-dependent lysine defatty-acylation may promote the localization of activated (GTP-loaded) K-Ras4a at endomembranes, which raises the possibility that lysine defatty-acylation may alter the signaling specificity of K-Ras4a by recruiting different effector proteins to endomembranes. We therefore investigated whether lysine defatty-acylation influenced the binding and activation of the three most well characterized Ras effectors: Raf1, PI3K, and RalGDS^61^. Co-immunoprecipitation demonstrated that neither the 3KR mutation nor Sirt2 KD altered the binding of K-Ras4a-G12V with Raf1, PI3K, or RalGDS (Fig. S8a). We also assessed the capacity of K-Ras4a-G12V and -G12V-3KR in Ctrl and Sirt2 KD cells to activate Raf1, PI3K, and RalGDS signaling pathways using phosphorylated Erk, phosphorylated Akt, and phosphorylated Jnk as reporters, respectively. K-Ras4a-G12V and -G12V-3KR induced comparable levels of Erk activation, which was not affected by Sirt2 KD. Sirt2 KD resulted in a reduction of Akt and Jnk activation, but the effect was similar for both K-Ras4a-G12V and -G12V-3KR (Fig. S8b), suggesting other Sirt2 targets are important for Akt and Jnk activation. These results suggest that SIRT2 catalyzed lysine defatty-acylation of K-Ras4a does not affect the activation of Raf1, PI3K or RalGDS by K-Ras4a.

To identify proteins whose binding to K-Ras4a is regulated by lysine fatty acylation, we performed a protein interactome study using stable isotope labeling by amino acids in cell culture (SILAC) (Fig. S9a). We cultured NIH3T3 cells with stable K-Ras4a-G12V and K-Ras4a-G12V-3KR overexpression in light-isotope- and heavy-isotope-labeled medium, respectively. We then performed FLAG IP, mixed the eluted fractions from both IPs, digested with trypsin and analyzed by MS to identify proteins with Heavy/Light (H/L) ratios > 1.3 or < 0.77, which were candidates that would potentially bind to K-Ras4a-G12V and K-Ras4a-G12V-3KR differently. The experiment was also repeated after swapping the heavy and light SILAC labels. Additionally, to confirm that the effect of the 3KR mutation on the K-Ras4a-G12V interactome was due to the lack of lysine fatty acylation, we also examined the K-Ras4a-G12V interactome in Ctrl and Sirt2 KD cells with SILAC, which enabled the identification of proteins (H/L > 1.3 or < 0.77) whose binding to K-Ras4a-G12V was regulated by Sirt2. Integration of the three interactome experiments resulted in 175 interacting proteins with at least two unique peptides and H/L ratio. Among them, nine proteins exhibited increased binding to K-Ras4a-G12V-3KR compared to K-Ras4a-G12V, and their interaction with K-Ras4a-G12V was inhibited by Sirt2 KD, suggesting that lysine defatty-acylation enhanced K-Ras4a-G12V interaction with these proteins. On the other hand, one protein showed decreased binding to K-Ras4a-G12V-3KR compared to K-Ras4a-G12V, and its interaction with K-Ras4a-G12V was increased by Sirt2 KD, suggesting that lysine defatty-acylation repressed K-Ras4a-G12V interaction with it (Fig. S9b).

Among these 10 proteins, the serine/threonine-protein kinase A-Raf and Apoptosis-inducing factor 1 (Aif), whose interaction with K-Ras4a-G12V might be increased by lysine defatty-acylation, attracted our attention. A-Raf is a member of the Raf family of serine/threonine-specific protein kinases, acts as a Ras effector and plays an important role in apoptosis ^62,63^ and tumorigenesis ^64–66^. In response to apoptotic stimuli, Aif is released from the mitochondrial intermembrane space into the cytosol and nucleus, where it functions as a proapoptotic factor^67^. Since suppression of apoptosis is linked to Ras-induced transformation^68^, it is plausible that A-Raf and Aif are involved in the regulation of K-Ras4a transformation activity by lysine fatty acylation. To test this hypothesis, we first validated the interactome results by co-IP. Although more interaction of Aif with K-Ras4a-G12V-3KR was observed than with K-Ras4a-G12V, Sirt2 KD did not affect the interaction of Aif with either K-Ras4a-G12V or K-Ras4a-G12V-3KR (Fig. S9c & d) and was not investigated further. However, a greater interaction of A-Raf with K-Ras4a-G12V-3KR was observed than with K-Ras4a-G12V, and Sirt2 KD significantly decreased the interaction of A-Raf with K-Ras4a-G12V but not with K-Ras4a-G12V-3KR (Fig. 6a & b). Thus, we concluded that A-Raf was an effector protein of K-Ras4a that was regulated by lysine fatty acylation and SIRT2. Unlike the effect of Sirt2 KD on K-Ras4a-G12V-A-Raf interaction, Sirt2 KD did not alter H-Ras-G12V-A-Raf interaction (Fig. 6a & b). As mentioned earlier, lysine fatty acylation did not affect the binding between C-Raf (Raf1) and K-Ras4a-G12V (Fig. S8a). We also assessed the interaction of K-Ras4a-G12V with another Raf family member, B-Raf. Co-IP indicated that neither 3KR mutation nor Sirt2 KD altered the binding of B-Raf to K-Ras4a-G12V (Fig. S8c). These results collectively demonstrate that removal of lysine fatty acylation from K-Ras4a by SIRT2 results in its preferential association with A-Raf, but not B-Raf or C-Raf.

Our results suggest that SIRT2-mediated lysine defatty-acylation does not affect the magnitude of K-Ras4a activation but promotes endomembrane localization of active K-Ras4a. It has been reported that the efficient activation of certain effector pathways by Ras is dependent on the entry of Ras to the endosomal compartment ^42,69^. Therefore, it is plausible that lysine defatty-acylation may facilitate the endomembrane recruitment of A-Raf by K-Ras4a, thereby increasing K-Ras4a oncogenic activity. Live cell imaging revealed that A-Raf colocalized with K-Ras4a-G12V at both the PM and endomembranes. K-Ras4a-G12V-3KR showed more colocalization with A-Raf on the endomembranes than K-Ras4a-G12V did. Sirt2 KD inhibited the endomembrane recruitment of A-Raf by K-Ras4a-G12V but not that by K-Ras4a-G12V-3KR (Fig. 6c & d). These results are in line with our hypothesis. Thus, it is likely that by regulating endomembrane recruitment of A-Raf, K-Ras4a lysine fatty acylation may alter its signaling output through A-Raf, thereby modulating its transforming activity.

**Figure 6.**
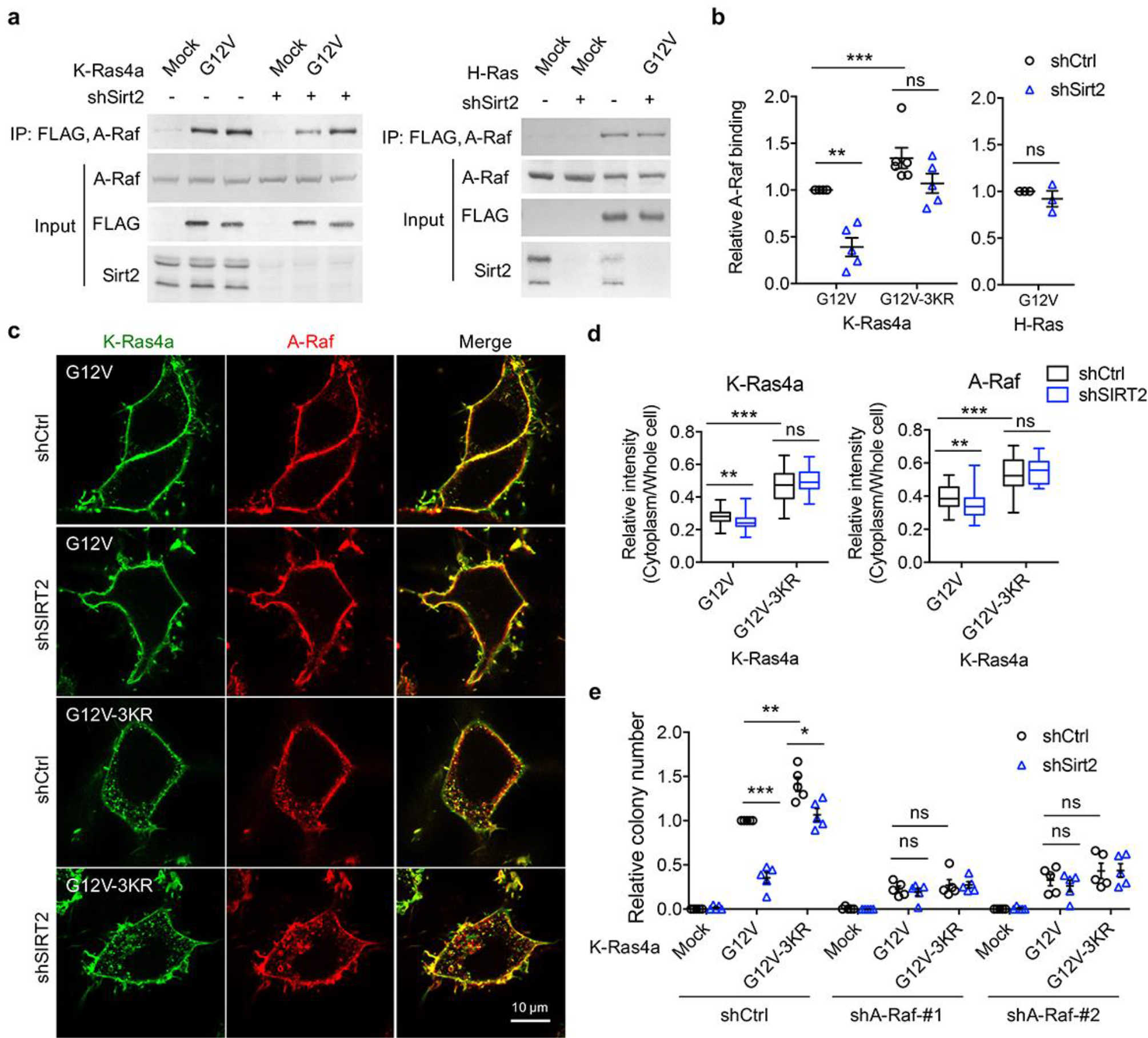
A-Raf is involved in the regulation of K-Ras4a transforming activity by lysine fatty acylation. (**a**) Co-IP of A-Raf with an anti-FLAG antibody in NIH 3T3 cells stably expressing Mock, FLAG-K-Ras4a-G12V, FLAG-K-Ras4a-G12V-3KR, or FLAG-H-Ras-G12V with Ctrl or Sirt2 KD. (**b**) Quantification of relative A-Raf binding levels in (a). The A-raf binding levels in in Ctrl KD cells were set to 1. Quantification was done with Fiji software. Signal intensity of A-Raf was normalized with the corresponding FLAG intensity. (**c**) Images showing the localization of GFP-K-Ras4a-G12V or -G12V-3KR and DsRed-A-Raf in live HEK293T cells with Ctrl or SIRT2 KD (by shSIRT2-#2). (**d**) Statistical analyses of the relative cytoplasm to whole cell intensity of K-Ras4a and A-Raf from (c) (n = 17 for all samples). (**e**) Anchorage-independent growth of NIH 3T3 cells stably expressing Mock, K-Ras4a-G12V or -G12V-3KR with Ctrl or Sirt2 KD, and Ctrl or A-Raf KDs. The y axis represents colony numbers relative to that of the Sirt2 or A-Raf Ctrl KD cells expressing K-Ras4a-G12V. Statistical evaluation in (b) and (e) was by unpaired two-tailed Student’s t test. Error bars represent SEM in at least three biological replicates or as indicated. Statistical evaluation in (d) was by two-way ANOVA. *Centre line* of the box plot represents the mean value, box represents the 95 *%* confidence interval, and whiskers represent the range of the values. \**P* < 0.05; \*\**P* < 0.01; \*\*\**P* < 0.001; ns, not significant. Representative images are shown.

While the functions of B-Raf and C-Raf in Ras-mediated oncogenic transformation have been well elucidated, the role of A-Raf in this process remains obscure^70^. So we next examined whether A-Raf plays a role in K-Ras4a-G12V mediated transformation using the soft agar colony formation assay. Inhibition of A-Raf expression with shRNA (Fig. S10a) partially suppressed K-Ras4a-G12V-induced colony formation, indicating that A-Raf is important for K-Ras4a-G12V mediated transformation. Moreover, A-Raf KD abrogated the 3KR mutation-mediated increase and Sirt2 KD-mediated decrease in the transformation activity of K-Ras4a-G12V (Fig. 6e & Fig. S10b), suggesting that A-Raf is important for the regulation of K-Ras4a transforming activity by SIRT2-dependent lysine defatty-acylation. These results further support the model that lysine defatty-acylation by SIRT2 enhances the recruitment of A-Raf to K-Ras4a at endomembranes, thereby promoting oncogenic activity of K-Ras4a.

## Discussion

Protein lysine fatty acylation was discovered over two decades ago ^4–8^. However, very little is known about its functional significance. Our current study furnishes a model where K-Ras4a is fatty acylated on lysine residues at its C-terminal HVR, and the removal of lysine fatty acylation by SIRT2 facilitates its endomembrane localization and interaction with A-Raf, thus enhancing its transforming activity (Fig. S11). These findings demonstrate that a Ras protein is modified and regulated by a previously underappreciated PTM, lysine fatty acylation, which expands not only the regulatory scheme for Ras proteins, but also the biological significance of lysine fatty acylation. Moreover, our study reveals the first lysine defatty-acylation substrate for SIRT2 and uncovers the physiological relevance of SIRT2 as a lysine defatty-acylase^12,14,15^.

We found that H-Ras and K-Ras4a possess lysine fatty acylation that could be hydrolyzed by sirtuins *in vitro* or in cells (Fig. 1, 2 & S2a). Although our attempt to detect N-Ras lysine fatty acylation by MS was not successful, the N-Ras-K169/170R (2KR) mutant presented decreased NH_2_OH-resisitant fatty acylation compared with WT, suggesting that N-Ras might be lysine fatty acylated (Fig. 1c & S1b). While cysteine palmitoylation of Ras proteins was discovered almost three decades ago^71^, lysine fatty acylation of Ras was not identified for several reasons. First, lysine fatty acylation did not emerge as a physiologically significant modification until recent years. Correspondingly, the possibility of lysine fatty acylation on Ras proteins had not been investigated previously. Second, previously people only focused on Ras cysteine palmitoylation because mutations of the palmitoylated cysteine to serine abolished the palmitoylation of H-Ras, N-Ras^72^ and K-Ras4a^19^ based on ^3^H-palmitic acid labeling. Therefore, lysine fatty acylation of Ras proteins might have been missed based on the mutagenesis results. Similar to, but slightly different from these previous reports, we found that mutating the palmitoylated cysteine of K-Ras4a decreased lysine fatty acylation by nearly 90% (Fig. 3c) but not completely. A similar effect of the palmitoylated cysteine to serine mutation was also observed for R-Ras2^25^. Last, although the palmitoylation sites for Ras proteins were characterized with chemoproteomic approaches based on acyl-biotin exchange (ABE) ^23,73^ or acyl-resin-assisted capture (acyl-RAC)^3,74^, these approaches are cysteine-centric and do not allow identification of amide-linked fatty acylation. Direct site identification of palmitoylation has been challenging owing to the low abundance and high hydrophobicity of modified peptides, which are easily lost during sample preparation^1^. Our current study highlights the regulation of Ras proteins by lysine fatty acylation and suggests that additional studies are required to understand the regulation of this important class of proteins. Many Ras superfamily of small GTPases contain lysine-rich sequences at their C-termini. It is therefore of great interest to us to investigate whether lysine fatty acylation could act as a general regulatory mechanism for many Ras-related small GTPases.

The discovery of lysine fatty acylation on K-Ras4a raises the question of the relative abundance of lysine versus cysteine fatty acylation. Semi-quantification of the fluorescence intensity from Alk14 labeling results enables us to roughly estimate the stoichiometry of lysine fatty acylation. Based on this, K-Ras4a exhibits nearly 50% of lysine fatty acylation relative to total fatty acylation (Fig. 1c). Therefore, the ratio of cysteine palmitoylation to lysine fatty acylation may be close to 1:1 on K-Ras4a. The 3KR mutation decreased K-Ras4a lysine fatty acylation by about 50% (Fig. 1f & 3e), suggesting that the C-terminal lysine fatty acylation regulated by SIRT2 is around 50% of the lysine fatty acylation and 25% of the total fatty acylation. Regarding endogenous K-Ras4a, by quantifying the K-Ras4a western blot signal from the streptavidin beads and supernatant in Fig. 2i, we estimated that about 28% and 50% of the fatty acylated K-Ras4a is lysine fatty acylated in Ctrl KD and SIRT2 KD HCT116 cells, respectively. Unfortunately, precise quantitation of protein fatty acylation still remains a significant unsolved challenge and we could not determine the ratio of fatty acylated versus unmodified K-Ras4a.

To study the physiological function of K-Ras4a lysine fatty acylation, we utilized the K-Ras4a-3KR mutant in combination with SIRT2 KD. The lysine-to-arginine mutant maintains the positive charge of the polybasic patch, which makes it a good lysine fatty acylation-deficient mimic. Recently, Zhou *et al.* reported that lysine and arginine residues are not equivalent in determining the membrane lipid binding specificity of K-Ras4b C-terminus, which raises the possibility that the effect of 3KR mutation might not be solely due to lack of lysine fatty acylation^75^. Likewise, changes in the SIRT2 KD cells could be mediated through other substrates for SIRT2. Therefore, it is critical to employ both the 3KR mutant and SIRT2 KD to rule out these possibilities. SIRT2 KD enhances the lysine fatty acylation of K-Ras4a WT but not the 3KR mutant. Thus, if a biological effect is due to lysine fatty acylation, SIRT2 should have a greater impact on the effect of K-Ras4a WT than that of the 3KR mutant. Of note, H-Ras, which shares similar properties with K-Ras4a, but is not a lysine defatty-acylase target for SIRT2, also serves as a good control for the effect of SIRT2 KD. Indeed, SIRT2 KD repressed the endomembrane localization, transforming activity, and A-Raf binding of K-Ras4a WT more than that of K-Ras4a 3KR or H-Ras, indicating that SIRT2-dependent lysine defatty-acylation facilitates endomembrane localization of K-Ras4a, enhances its interaction with A-Raf, and thus promotes cellular transformation.

Goodwin *et al.* previously reported that cysteine depalmitoylated H-Ras and N-Ras traffic to and from the Golgi complex by a nonvesicular mechanism, and suggested a model where cysteine palmitoylation traps Ras on membranes, enabling Ras to undergo vesicular transport^76^. In line with this, Tsai *et al.*^19^ and we observed that the K-Ras4a-C180S mutant, which possesses no cysteine palmitoylation and little lysine fatty acylation, localizes to ER/Golgi-like internal membranes (Fig. S5e & f). Differently, we found that removal of the lysine fatty acylation by SIRT2, which results in K-Ras4a with only cysteine palmitoylation, promotes endomembrane localization of K-Ras4a (Fig. 4a). This evidence supports the model that cysteine palmitoylation enables K-Ras4a to undergo vesicular transport, whereas lysine fatty acylation blocks K-Ras4a translocation from the PM to endomembranes (Fig. S11). Furthermore, the C180S mutant suppressed K-Ras4a-G12V-mediated anchorage-independent growth and activation of MAPK signaling^19,20^. In contrast, the 3KR mutant increased K-Ras4a-G12V-mediated anchorage-independent growth (Fig. 5b & c), exhibited no effect on MAPK signaling (Fig. S8), but activated A-Raf instead (Fig. 6a). Considering the possibility that lysine fatty acylation largely relies on cysteine palmitoylation to occur, it is likely that the reversible lysine fatty acylation adds a layer of regulation for K-Ras4a above that of cysteine palmitoylation.

In the GTP-bound active form, Ras proteins bind directly to the Ras binding domain (RBD) of Raf, then form secondary interactions with a cysteine-rich domain (CRD). Although the RBD and CRD are highly conserved in all Raf isozymes, there is evidence for different binding affinities for Ras proteins to the individual Raf proteins^77,78^. For example, Weber *et al.* reported that A-Raf presents significantly lower binding affinities to H-Ras-G12V as compared to C-Raf because the Ras-binding interface of C-Raf differs from A-Raf by a conservative arginine to lysine exchange at residue 59 or 22 respectively^79^. Furthermore, Williams *et al.*^80^ and Fischer *et al.^81^* found that farnesylation of H-Ras is required for its binding to C-Raf but not to B-Raf, implying the involvement of Ras C-terminal PTM in regulating Ras-Raf interactions. Based on these previous studies, it is likely that the C-terminal PTM of K-Ras4a regulates its interaction with Raf isozymes and that lysine fatty acylation may inhibit the binding of K-Ras4a to A-Raf but not to B-Raf and C-Raf.

Mutations that activate Ras are found in about 30% of all human tumors screened. *KRAS* mutations, which affects both K-Ras4a and K-Ras4b, occur most frequently, accounting for 86% of *RAS*-driven cancers^82^. Though K-Ras4a is homologous to the transforming cDNA identified in Kirsten rat sarcoma virus^83^, its function and regulation is less characterized compared to K-Ras4b. Recent studies showed that K-Ras4a is widely expressed in human cancers, suggesting that K-Ras4a plays a significant role in KRAS-driven tumors^19,20^. Our findings reveal that K-Ras4a is regulated by SIRT2-dependent lysine defatty-acylation. Depletion of SIRT2 increased lysine fatty acylation and diminished oncogenic transforming activity of K-Ras4a, suggesting that interference with K-Ras4a lysine fatty acylation could be an approach to anti-K-Ras therapy.

The seven mammalian sirtuins, SIRT1-7, are implicated in various biological pathways and are considered potential targets against a number of human diseases^55^. So far, the known biological functions of sirtuins have been mainly attributed to their deacetylase activities. Although sirtuins are increasingly recognized as lysine deacylases in addition to deacetylases, the biological significance of sirtuins as lysine deacylases remains largely unknown^84^. Our work here identifies the first physiological defatty-acylation substrate for SIRT2. Since protein acyl lysine modifications likely use acyl-CoA molecules as the acyl donors, the cellular metabolic state can affect acyl lysine PTMs by altering the concentration of acyl-CoA molecules. Sirtuins requires NAD as a co-substrate and the NAD level is regulated by cellular metabolism. Thus SIRT2 may provide an additional link between K-Ras4a signaling and cellular metabolism. Given that Ras proteins play critical roles in many human cancers, SIRT2, as a Ras regulator, may be an important therapeutic target for cancer, which is consistent with several recent reports ^50,53, 54,57,58,85–87^. The physiological and pathophysiological roles of SIRT2 thus merit further investigation.

## Acknowledgements

This work is supported in part by a grant (1R01GM121540-01A1) from NIH. H.J. is a Howard Hughes Medical Institute International Student Research Fellow. We thank Cornell University Biotechnology Resource Center (BRC) Imaging Facility for the support on the confocal microscopy usage.

## Author Contributions

H.J. designed and performed all the biochemical and cellular studies except those noted below. X.Z. synthesized K-Ras4a-C180myr peptide, carried out MS analyses of H-Ras and K-Ras4a lysine fatty acylation, *in vitro* cysteine depalmitoylation assay, ^32^P-NAD assay, and K-Ras4a intereactome study.S.A.W. purified SIRT2 protein and performed the Alk14 labeling experiment for the K-Ras4a single KR mutants, C186S and C180S mutants. X.C. validated H.J.’s results on cell proliferation and soft agar colony formation. N.A.S. synthesized the Alk14 probe. M.E.L provided pCMV5-*HRAS*, pCMV5-*NRAS*, pCMV5-*K-RAS4B* and pCMV5-*RalA* plasmids and consultation on Ras GTPases. H.L. directed and supervised all the studies. H.J., X.Z. and H.L. wrote the manuscript and all authors reviewed and approved the manuscript.

## Methods

### Reagents, antibodies and plasmids

Chemicals from commercial sources were obtained in the highest purity available. Alk14, Rhodamin-N_3_ and Biotin-N_3_ were synthesized as previously reported^88^. Trichostatin A (TSA, T8552), protease inhibitor cocktail (P8340), phosphatase inhibitor cocktail (P0044), Azide-PEG3-biotin (762024), Tris[(1-benzyl-1H-1,2,3-triazol-4-yl)methyl]amine (TBTA, 678937), Tris(2-carboxyethyl)phosphine (TCEP, 75259), hydroxylamine (NH_2_OH, 159417), NAD (NAD100-RO), Puromycin (P8833), Crystal Violet (C0775), low-melting point agarose (A0701), triple FLAG peptide (F4799), L-lysine (L9037), L-arginine (A8094), [^13^C_6_, ^15^N_2_]-L-lysine (608041) and [^13^C_6_,^15^N_4_]-L-arginine (608033) were purchased from Sigma-Aldrich. The anti-human SIRT1 (05-1243), anti-RalA (ABS223) and anti-Ras (Y13-259, OP01A) antibodies were from EMD Chemicals Inc. The anti-SIRT2 (ab134171), Transferrin Receptor (ab84036) antibodies were from Abcam. The anti-SIRT2 (12650), Phospho-Erk1/2 (Thr202/204) (9101), Erk1/2 (4696), Phospho-Akt (Thr308) (9275), Phospho-Akt (Ser473) (9271), Akt (4691), Phospho-SAPK/JNK (Thr182/Tyr185) (4668), SAPK/JNK (9252), Syntaxin 6 (STX6, 2869), EEA1 (3288) antibodies were purchased from Cell Signaling Technology. The anti-β-Actin (sc-4777), K-Ras4a (sc-522), A-Raf (sc-408), B-Raf (sc-166), C-Raf (sc-227), Na/K-ATPase (sc-21712), GAPDH (sc-47724), the normal rat IgG (sc-2026) and the goat antimouse/rabbit/Rat IgG-horseradish peroxidase-conjugated antibodies were purchased from Santa Cruz Biotechnology. The anti-Acetyl Lysine antibody (ICP0380) was obtained from Immunechem. The antiFLAG M2 antibody conjugated with horseradish peroxidase (A8592) and the anti-FLAG M2 affinity gel (A2220) were from Sigma-Aldrich. Enzyme-linked chemiluminescence (ECL) plus (32132) western blotting detection reagent, Cy3-conjugated goat anti-rabbit IgG (H+L) secondary antibody (A10520) and the high capacity Streptavidin agarose (20357) were purchased from Thermo Fisher Scientific. FuGene 6 (E2692) transfection reagent and sequencing grade modified trypsin (V5111) were purchased from Promega. ^32^P-NAD^+^ was purchased from PerkinElmer. Saponin (S0019-25G) was from TCI America. Sep-Pak C18 cartridge was purchased from Waters.

The pLKO.1-puro lentiviral shRNAs constructs for luciferase and human SIRT1, SIRT2, and mouse SIRT2 were purchased from Sigma-Aldrich. Luciferase shRNA (SHC007), human SIRT1 shRNAs (#1, TRCN0000018980, #2, TRCN0000018981), human SIRT2 shRNAs (#1, TRCN0000040219, #2, TRCN0000310335), mouse SIRT2 shRNA (TRCN0000012118), mouse A-Raf shRNAs (#1, TRCN0000022612, #2, TRCN0000022610) were used. The human *K-RAS4A* expression vector with N-terminal FLAG tag was obtained by RT-PCR amplification of *K-RAS4A* and subcloning *via* EcoRI and SalI sites into pCMV5 vector. The human *K-RAS4A* lentiviral vector was obtained by inserting FLAG-*K-RAS4A* into pCDH-CMV-MCS-EF1-Puro vector between the EcoRI and NotI sites. The human *H-RAS* lentiviral vector was obtained by inserting FLAG-*HRAS* into pCDH-CMV-MCS-EF1-Puro vector between the EcoRI and BamHI sites. The GFP-K-Ras4a and GFP-H-Ras expression vectors were constructed by inserting *K-RAS4A and H-RAS* cDNA into pGFP1-C1 vector between the BglII and SalI sites, respectively. The human *STX6* expression vector with N-terminal FLAG tag was constructed by inserting FLAG-STX6 cDNA into pCMV-tag-4a vector between the EcoRI and XhoI sites. To generate the expression vector for human SIRT2 with C-terminal FLAG tag, full-length human *SIRT2* cDNA was amplified by PCR and inserted into pCMV-tag-4a vector between the BamHI and XhoI sites. The expression vectors for H-Ras, N-Ras, K-RAS4A mutants and SIRT2-H187Y were generated by QuikChange site-directed mutagenesis^89^. The DsRed cDNA without stop codon was inserted using NotI and BamHI sites into pCMV-tag-4a to generate pCMV-tag-4a-DsRed-C vector that enables cloning of gene of interest with N-terminal DsRed. The DsRed-RBD expression vector was constructed by inserting cDNA coding the Ras-binding domain of human Raf1 (aa51-131) into pCMV-tag-4a-DsRed-C vector between EcoRV and XhoI sites. The DsRed-A-Raf expression vector was generated by inserting mouse *Araf* cDNA into pCMV-tag-4a-DsRed-C vector using BamHI and EcoRI sites. DsRed-GalT plasmid^90^ was obtained from Dr. Yuxin Mao (Cornell Univeristy, Ithaca, NY). Expression vectors for mCherry-Sec61 beta (Addgene plasmid #49155)^91^, mCherry-Rab11 (Addgene plasmid #55124), DsRed-Rab7 (Addgene plasmid #12661)^92^ and Lamp1-RFP (Addgene plasmid #1817)^93^ were gifts from Gia Voeltz, Michael Davidson, Richard Pagano and Walther Mothes, respectively.

### Cell culture, transfection and transduction

Human HEK293T cells were grown in DMEM media (11965-092, Gibco) with 10% heat inactivated (HI) fetal bovine serum (FBS, 26140079, Gibco). Mouse embryonic fibroblast NIH3T3 cells were grown in DMEM media supplemented with non-essential amino acids (11140050, Gibco) and 15% HI FBS. Human HCT116 cells were grown in McCoy’s 5A media (16600082, Gibco) with 10% HI FBS. The cell lines were purchased from American Type Culture Collection (ATCC). The cell lines were not further authenticated after purchase from ATCC. All cell lines were tested for and showed no mycoplasma contamination.

For SILAC experiments, ‘light’ NIH3T3 cells were maintained in DMEM media for SILAC (88420, Thermo Fisher Scientific) supplemented with 100 mg/L [^12^C_6_, ^14^N_2_]-L-lysine, 100 mg/L [^12^C_6_, ^14^N_4_]-L-arginine, non-essential amino acids, and 15% dialyzed FBS (26400036, Thermo Fisher Scientific); ‘heavy’ NIH3T3 cells were cultured in DMEM media for SILAC supplemented with 100 mg/L [^13^C_6_, ^15^N_2_]-L-lysine, 100 mg/L [^13^C_6_, ^15^N_4_]-L-arginine, non-essential amino acids, and 15% dialyzed FBS. Cells were cultured in SILAC media for at least six doubling times to achieve maximum incorporation of ‘labeled’ amino acids into proteins before the interactome study was performed.

To transiently overexpress proteins of interest in cells, the expression vectors were transfected into cells using FuGene 6 according to the manufacturer’s protocol. Empty vector was transfected as a negative control. Lentiviral infection for overexpressing H-Ras, K-Ras4a WT and mutants or knocking down SIRT1, SIRT2 and A-Raf was performed as previously described ^9,53^. Puromycin (3 μg/mL for NIH3T3 cells, 1.5 μg/mL for HEK293T cells) was added to the cell culture media to select NIH3T3 cells with stable overexpression of Mock (pCDH empty vector control), K-Ras4a-G12V, K-Ras4a-G12V-3KR or H-Ras-G12V as well as HEK293T cells with stable luciferase KD (Ctrl KD), SIRT1 KD, or SIRT2 KD.

### Immunoprecipitation of Alk14-labeled proteins of interest

HEK293T cells (parental cells, luciferase KD, SIRT1 KD, or SIRT2 KD) were transiently transfected to express FLAG-tagged protein of interest overnight. The cells were then cultured with fresh medium containing 50 μM Alk14 for 6 h. Cells were collected and lysed in 1% NP-40 lysis buffer (25 mM Tris-HCl pH 8.0, 150 mM NaCl, 10% glycerol, 1% Nonidet P-40) with protease inhibitor cocktail. The supernatant was collected after centrifugation at 16,000g for 20min at 4°C. Protein concentration was determined by Bradford assay (23200, Thermo Fisher Scientific). 0.5-1 mg cell lysate was incubated with 10μL suspension of antiFLAG M2 affinity gel for 2h at 4 °C. The affinity gel was then centrifuged at 500 *g* for 2 min at 4 °C, washed three times with 1 mL IP washing buffer (25 mM Tris-HCl pH 8.0, 150 mM NaCl, 0.2% Nonidet P-40) and used for further experiments.

### Detection of fatty acylation on protein of interest by on-beads click chemistry and in-gel fluorescence

The immunopurified protein with Alk14 labeling was suspended in 20 μL IP washing buffer for click chemistry. Rh-N_3_ (3 μL of 1 mM solution in DMF, final concentration 150 μM) was added to the above suspension, followed by the addition of TBTA (1 μL 10 mM solution in DMF, final concentration 500 pM), CuSO_4_ (1 μL of 40 mM solution in H_2_O, final concentration 2 mM), and TCEP (1 μL of 40 mM solution in H_2_O, final concentration 2 mM). The click chemistry reaction was allowed to proceed at room temperature for 30 min. The reaction mixture was mixed with 10 μL of 6 × protein loading buffer and heated at 95°C for 10 min. After centrifugation at 16,000 g for 2 min at room temperature, 15 μL of the supernatant was treated with NH_2_OH (pH 8.0, 1 μL of 5 M solution in H_2_O, final concentration 300 mM) or equivalent volume of water (negative control) at 95 °C for 7 min. The samples were resolved by SDS-PAGE. Rhodamine fluorescence signal was recorded by Typhoon 9400 Variable Mode Imager (GE Healthcare Life Sciences, Piscataway, NJ) with setting of Green (532nm)/580BP30 PMT 500 V (normal sensitivity). Fiji software^94^ was used for quantification of the fluorescence intensity. Signal intensity of in-gel fluorescence was normalized with respect to that of the corresponding FLAG western blot.

### Defatty-acylation of K-Ras4a by sirtuins *in vitro*

The *Plasmodium falciparum* Sir2A (PfSir2A)^26^, the human SIRT1^95^, SIRT2^53^, SIRT3^95^ and SIRT6^9^ were expressed as previously described. The immunoprecipated Ras with Alk14 labeling on anti-FLAG affinity gel was suspended in 25 μl of assay buffer (50 mM Tris-HCl, pH 8.0, 100 mM NaCl, 2 mM MgCl_2_, 1 mM DTT) with 10 μM of PfSir2A or 5 μM of SIRT1, SIRT2, SIRT3, SIRT6 or the corresponding amount of BSA and with or without 1 mM NAD and incubated at 37 °C for 30 min (SIRT2) or 1 h (PfSir2A, SIRT1, 3 and 6). The reaction was stopped by washing the affinity gel using 1 mL of IP washing buffer for 3 times. On-bead click chemistry and in-gel fluorescence were carried out as described above.

### High-performance liquid chromatography (HPLC)-based SIRT2 activity assay

SIRT2 or SIRT2-H187Y (1 μM) was incubated in 60 μL of reaction buffer (20 mM Tris, pH 8.0, 1 mM DTT, 1 mM NAD) with 32 μM acetyl H3K9, myristoyl H3K9, or myristoyl K-Ras4a-C180 peptides (Fig. S12), respectively, at 37°C for 10 min (deacetylation) or 20 min (demyristoylation). Reactions were quenched with 60 μL ice-cold acetonitrile and spun down at 18,000 g for 10 min to remove the precipitated protein. The supernatant was then analyzed by HPLC on a Kinetex XB-C18 column (100 A, 75 mm × 4.6 mm, 2.6 μm, Phenomenex). The peak areas were integrated and the conversion rate was calculated from the ratio of the free H3K9 peptide peak area over the total peak areas of the substrate and product peptides.

### Western blot

Western blot analysis was performed as described previously^9^. The proteins of interest were detected using ECL plus and visualized using the Typhoon 9400 Variable Mode Imager (GE Healthcare). Quantification of signal intensity from western blots was done using Fiji software. To assess the effect of lysine fatty acylation on the signaling output of K-Ras4a-G12V through Erk, Akt and Jnk, NIH3T3 cells stably expressing Mock, FLAG-K-Ras4a-G12V or -G12V-3KR were infected with lentivirus carrying luciferase (Ctrl) or mouse Sirt2 shRNA for 3 days, collected and lysed in 1% NP-40 lysis buffer with protease inhibitor cocktail and phosphatase inhibitor cocktail. Cell lysates were then subjected to western blot for the analyses of indicated proteins.

To detect acetyl lysine on K-Ras4a, HEK293T cells with stable Ctrl KD or SIRT2 KD were transfected with empty vector or pCMV5-K-*RAS4A* overnight. The cells were then treated with ethanol or trichostatin A (TSA, 1 μM) for 1 h. The cells were collected and lysed in 1% NP-40 lysis buffer with protease inhibitor cocktail. Cell lysates (~3 mg), with/without overexpression of K-Ras4a, were incubated with 10 μL of anti-FLAG M2 affinity gel suspension for 2h at 4°C. The affinity gel was washed three times with 1 mL of IP washing buffer and then heated in 15 μL of 2 × protein loading buffer at 95 °C for 10 min. The supernatant was then resolved by SDS-PAGE and the acetylation of K-Ras4a was examined by western blot using anti-acetyllysine antibody after transfer to a PVDF membrane. Total cell lysates from TSA-treated HEK293T cells were used as a positive control for the acetyllysine blot. After recording the acetyl-lysine signal, the PVDF membrane was stained with Coomassie blue to detect K-Ras4a protein. A western blot using anti-FLAG antibody was carried out in parallel to demonstrate equal loading of K-Ras4a.

### Subcellular fractionation

HEK293T cells were transfected with pCMV5-K-*RAS4a* and cultured for overnight before being collected. Cell pellets were re-suspended in subcellular fraction buffer (250 mM Sucrose, 20 mM HEPES, pH 7.4, 10 mM KCl, 1.5 mM MgCl_2_, 1 mM EDTA, 1 mM EGTA and 1 mM DTT) containing protease inhibitor cocktail and homogenized on ice by 10 passes through a 25-gauge syringe needle. Nuclei and intact cells were removed by centrifugation at 3,000 rpm for 5 min. The mitochondrial fraction was removed by centrifuging the postnuclear supernatant at 8,000 rpm for 5 min. The supernatant was ultracentrifuged at 40,000 rpm for 1 h. The resulting supernatant (cytosol fraction) was concentrated through the filter. The pellet (membrane fraction) was washed with subcellular fraction buffer, re-centrifuged for 45 min and dissolved in 4% SDS lysis buffer (4% SDS, 50 mM triethanolamine pH 7.4, and 150 mM NaCl). Equivalent portions of the cytosol and membrane fractions were then subjected to western blot analyses.

### Co-Immunoprecipitation

To examine the interaction between FLAG-tagged K-Ras4a and SIRT2, HEK293T cells transfected with empty vector or pCMV5-FLAG-K-*RAS4A* were cultured overnight, collected and lysed in 1% NP-40 lysis buffer with protease inhibitor cocktail. To examine the interaction between FLAG-tagged K-Ras4a-G12V/K-Ras4a-G12V-3KR or H-Ras-G12V and A-Raf/B-Raf/C-Raf (Raf1)/p110α/RalGDS, NIH3T3 cells stably expressing Mock, FLAG-K-Ras4a-G12V, -G12V-3KR, or FLAG-H-Ras-G12V were infected with lentivirus carrying luciferase (Ctrl) or mouse Sirt2 shRNA for 3 days, collected and lysed in 1% NP-40 lysis buffer with protease inhibitor cocktail. For both experiments, total cell lysates (2 mg of total protein for detecting SIRT1/2, 50 μg for A-Raf and c-Raf, 1 mg for B-Raf, p110α and RalGDS, determined by Bradford assay) were incubated with 10μL suspension of anti-FLAG M2 affinity gel for 2h at 4°C. The resulting affinity gel was washed three times with 1 mL IP washing buffer and heated in protein loading buffer (2 × final concentration) at 95 °C for 10 min. Western blot was then performed to detect levels of the indicated proteins.

### Detection of lysine fatty acylation on K-Ras4a using the ^32^P-NAD assay

The ^32^P-NAD assays were carried out as described previously with minor modification^30^. HEK293T cells were transfected with empty pCMV5 vector or pCMV5-K-*RAS4A* overnight and lysed in 1% NP-40 lysis buffer with protease inhibitor cocktail. For each reaction, cell lysates (3 mg of total protein, determined by Bradford assay) were incubated with 10μL suspension of anti-FLAG M2 affinity gel for 2 h at 4 °C. The affinity gel was washed three times with 1 mL of IP washing buffer. The resulting anti-FLAG affinity gel or the synthetic acetyl and myristoyl H3K9 peptides^30^ (25 μM, positive control) were mixed with 10 μL solutions containing 1 μCi ^32^P-NAD, 50 mM Tris-HCl pH 8.0, 150 mM NaCl, 1 mM DTT. The reactions were incubated with 1 μM BSA (negative control), SIRT2, or SIRT2-H187Y at 37°C for 30 min. A total of 2 μL of each reaction were spotted onto silica gel TLC plates and developed with 7:3 ethanol:ammonium bicarbonate (1 M aqueous solution). After development, the plates were air-dried and exposed to a PhosphorImaging screen (GE Healthcare). The signal was detected using Typhoon 9400 Variable Mode Imager (GE Healthcare).

### Biotin pull-down of lysine fatty acylated endogenous K-Ras4a

The assay was carried out as previously described with some modifications ^31^. Briefly, HCT116 cells were infected with lentivirus carrying luciferase (Ctrl) or SIRT2 shRNA for 3 days and treated without or with Alk14 (50 μM) for 6 h before being collected. Total proteins were then extracted using 1% NP-40 lysis buffer with protease inhibitor cocktail. 10 mg of total protein extract was subjected to click reaction with 100 μM Biotin-N_3_, 500 μM TBTA, 1 mM CuSO_4_ and 1 mM TCEP in a final volume of 5 mL. The reaction was allowed to proceed at room temperature for 1 h. Proteins were precipitated by adding 4 volumes of ice-cold methanol, 3 volumes of water, and 1.5 volumes of chloroform. Precipitated proteins were pelleted by centrifugation (4,500× g, 20 min, 4 °C), washed twice with 50 mL of ice-cold methanol and air-dried. The protein pellet was suspended in 4% SDS buffer (4% SDS, 50 mM triethanolamine pH 7.4, and 150 mM NaCl, 10 mM EDTA). The solubilized protein mixture was diluted to 1% SDS with 1% Brij 97 (in 50 mM triethanolamine pH 7.4, and 150 mM NaCl) and incubated with streptavidin agarose (0.2 ml slurry for 1 mg of protein) for 1 h at room temperature. The streptavidin beads were washed three times with 10 mL of 1% SDS in PBS buffer. The streptavidin beads were incubated with 1 M NH_2_OH (pH 8.0) in 300 μL of 1% SDS PBS buffer for 1 h at room temperature to elute proteins with only cysteine fatty acylation. The resulting supernatant was concentrated to 20 μL final volume using the Amicon Ultra-0.5 Centrifugal Filter (UFC501008, EMD Millipore). The resulting streptavidin beads were washed three times with 1% SDS in PBS buffer. Both the concentrated supernatant and washed beads were heated in protein loading buffer (2 × final concentration) at 95°C for 10 min and subjected to western blot analyses.

### Confocal microscopy

Cells were seeded in 35-mm glass bottom dishes (MatTek) and transfected with relevant constructs overnight.

For live cell imaging, cells were incubated in the Live Cell Imaging Solution (A14291DJ, Thermo Fisher Scientific) and imaged with a Zeiss 880 confocal/multiphoton inverted microscope (Carl Zeiss MicroImaging, Inc., Thornwood, NY) in a humidified metabolic chamber maintained at 37 °C and 5% CO2. For time-lapse movies, 60 single section images were recorded at 1 sec intervals for 1 min.

For immunofluorescence, cells were rinsed with 1 × PBS twice and fixed with 4% paraformaldehyde (v/v in 1 × PBS) for 15 min. The fixed cells were washed twice with 1 × PBS, permeabilized and blocked with 0.1% Saponin/5% BSA/1 × PBS for 30 min. The cells were then incubated overnight at 4 °C in dark with indicated primary antibody at 1/50 - 1/100 dilution (in 0.1% Saponin/5% BSA/1 × PBS). Cells were washed with 0.1% Saponin/1 × PBS three times and incubated with Cy3-conjugated goat anti-rabbit IgG (H+L) secondary antibody at 1/1000 dilution (in 0.1% Saponin/5% BSA/1 × PBS) at room temperature in dark for 1 h. Samples were washed with 0.1% Saponin/1 × PBS three times and mounted with Fluoromount-G^®^ (0100-01) from SouthernBiotech before imaging with Zeiss LSM880 inverted confocal microscopy. Images were processed with Fiji software.

For colocalization analyses of GFP-K-Ras4a WT or -3KR with various intracellular membrane markers, live cell imaging was performed for colocalization with mCherry-Sec61, DsRed-GalT, mCherry-Rab11, DsRed-Rab7 and Lamp1-RFP; immunofluorescence was performed for colocalization with STX6 (1/50 dilution for anti-STX6 antibody) and EEA1 (1/100 dilution for anti-EEA1 antibody).

### Quantitative analyses of colocalization and fluorescence intensity

Fiji software was used for quantification. To quantify the degree of cytoplasmic colocalization, background was subtracted, then the cytoplasm area was selected and quantified for each cell examined. Pearson’s correlation coefficient^96^ was calculated using Fiji plug-in Coloc2 program (http://fiji.sc/Coloc_2) on a single plane between the two indicated fluorescent signals. To quantify fluorescence intensity, background was subtracted and the cytoplasm area or the whole cell was selected for integrated signal intensity quantification. Relative cytoplasm with respect to whole cell fluorescence intensity was presented.

### Soft agar colony formation assay

To assess the effect of Ctrl or Sirt2 KD on K-Ras4a-G12V, - G12V-3KR or H-Ras-G12V-mediated anchorage-independent growth, NIH3T3 cells with stable overexpression of Mock, K-Ras4a-G12V or -G12V-3KR were infected with Ctrl shRNA- or Sirt2 shRNA-carrying lentivirus for 6 h and cultured in complete medium for another 72 h before being seeded for soft agar colony formation assay.

To determine the effect of Ctrl or A-Raf KD, NIH3T3 cells with stable overexpression of Mock (pCDH empty vector control), K-Ras4a-G12V or -G12V-3KR were first infected with Ctrl shRNA- or Sirt2 shRNA-carrying lentivirus for 6 h and then with Ctrl shRNA- or A-Raf shRNAs for another 6 h. The infected cells were then cultured in complete medium for another 72 h before being seeded for soft agar colony formation assay.

0.6% base low-melting point agarose (LMP) and 0.3% top LMP were prepared by mixing 1.2% LMP in H_2_O and 0.6% LMP in H_2_O, respectively, with 2 × complete medium in 1:1 (v/v) ratio. 1.5 mL of 0.6% base LMP was added to each well of 6-well plate and allowed to solidify for 30 min at room temperature. Then 5.0 ×10^3^ cells were resuspended in 0.3% LMP top LMP and plated onto 6-well plate pre-coated with the base LMP. 150 μL of complete medium was added on top of the 0.3% LMP and refreshed every 3 days. After 14 days of culture, colonies were stained with 0.1% crystal violet (m/v in 25% methanol) for 30 min, rinsed with 50% methanol, and counted.

### Cell proliferation assay

NIH3T3 cells with stable overexpression of Mock, K-Ras4a-G12V, or -G12V-3KR were seeded in 12-well plate at a density of 1.5 × 10^4^ cells/well 24 h before being infected with luciferase (Ctrl) shRNA- or Sirt2 shRNA-carrying lentivirus for 0 or 5 days. After knocking down Sirt2 for the indicated time, cells were washed with 1 × PBS, fixed with ice-cold methanol for 10 min and then stained with 0.25% crystal violet (m/v, in 25% methanol) for 10 min. The stained cells were washed with running distilled water, air-dried and solubilized in 200-800 μL of 0.5% SDS in 50% ethanol. Absorbance of the resulting solution was measured at 550 nm.

### Transwell migration assay

NIH3T3 cells with stable overexpression of Mock, K-Ras4a-G12V or -G12V-3KR were infected with Ctrl shRNA- or Sirt2 shRNA-carrying lentivirus for 6 h and cultured in complete medium for another 72 h. Cells were cultured in serum-free medium for 12 h before the assay. The assay was performed in 24-well Transwell plate with 8 mm polycarbonate sterile membrane (Corning Incorporated). Cells were plated in the upper chamber (20,000 cells/insert) in 200 μL of serum-free medium. Inserts were then placed in wells containing 600 μL of medium supplemented with 10% FBS. 12 h later, cells on the upper surface of the filter were detached with a cotton swab and cells on the lower surface of the filters were fixed with ice-cold methanol for 10 min and stained with 0.1% crystal violet for 15 min. The cells were then rinsed with distilled water, photographed and counted. Migration was quantified by counting the migrated cells in ten random microscopic fields.

### Active Ras pull-down and detection

Ras activity was determined using a Ras binding domain of Raf1 (RBD) pull-down assay kit (16117, Thermo Fisher Scientific) by following the manufacturer's instructions. Briefly, to determine RBD binding of K-Ras4a WT and -3KR in Ctrl or SIRT2 KD cells, HEK293T cells expressing FLAG-K-Ras4a WT or 3KR were infected with luciferase (Ctrl) shRNA- or human SIRT2 shRNA-carrying lentivirus for 6 h and cultured in complete medium for another 72 h. Cells were then serum-starved overnight and treated with 100 ng/mL EGF for 0, 5 and 15 min. At the end of treatment, cells were rinsed with ice-cold 1 × PBS and scraped on ice in lysis buffer containing 25mM Tris pH 7.2, 150 mM NaCl, 5 mM MgCl_2_, 1% NP-40 and 5% glycerol and 1 × protease inhibitor cocktail. The samples were collected, vortexed, incubated on ice for 5 min and centrifuged at 16,000 g at 4 °C for 15 min to remove cellular debris. Protein concentration was measured by Bradford assay. Equal amounts of lysate (500 μg) were incubated with RBD-coated agarose beads at 4 °C for 1 h. The beads were then washed three times with ice-cold lysis buffer, boiled for 5 min at 95 °C, and active Ras was analysed by western blot using Ras-specific antibodies (16117, Thermo Scientific). For comparison to total Ras protein, 1% of total lysates used for pull-down was analysed by immunoblot.

To determine RBD binding of K-Ras4a-G12V and -G12V-3KR in Ctrl or SIRT2 KD cells, HEK293T cells were transfected with pCMV5-FLAG-K-Ras4a-G12V or -G12V-3KR and infected with luciferase (Ctrl) shRNA- or human SIRT2 shRNA-carrying lentivirus 12 h after the transfection. 72 h later, cells were cultured in FBS-free or complete medium for another 12 h before being subjected to RBD pull-down as described above.

### K-Ras4a interactome by SILAC

Three SILAC experiments were performed to determine K-Ras4a-G12V or -G12V-3KR interacting proteins: (1) NIH3T3 cells stably overexpressing FLAG-K-Ras4a-G12V cultured in DMEM with [^12^C_6_, ^14^N_2_]-L-lysine and [^12^C_6_, ^14^N_4_]-L-arginine as “light” cells, and NIH3T3 cells stably overexpressing FLAG-K-Ras4a-G12V-3KR cultured in DMEM with [^13^C_6_, ^15^N_2_]-L-lysine and [^13^C_6_, ^15^N_4_]-L-arginine as “heavy” cells. (2) NIH3T3 cells stably overexpressing FLAG-K-Ras4a-G12V-3KR cultured in DMEM with [^12^C_6_, ^14^N_2_]-L-lysine and [^12^C_6_, ^14^N_4_]-L-arginine as “light” cells, and NIH3T3 cells stably overexpressing FLAG-K-Ras4a-G12V cultured in DMEM with[^13^C_6_, ^15^N_2_]-L-lysine and [^13^C_6_, ^15^N_4_]-L-arginine as “heavy” cells. The second group served as the reverse SILAC of the first group. (3) NIH3T3 cells stably overexpressing FLAG-K-Ras4a-G12V and transiently transduced with luciferase (Ctrl) shRNA cultured in DMEM with [^13^C_6_, ^15^N_2_]-L-lysine and [^13^C_6_, ^15^N_4_]-L-arginine as “heavy” cells, and NIH3T3 cells stably overexpressing FLAG-K-Ras4a-G12V and transiently transduced with mouse Sirt2 shRNA cultured in DMEM with [^12^C_6_, ^14^N_2_]-L-lysine and [^12^C_6_, ^14^N_4_]-L-arginine as “light” cells.

Cells were collected and lysed in 1% NP-40 lysis buffer containing protease inhibitor cocktail. Protein concentration was quantified by Bradford assay, and 8 mg of total protein from each sample was subjected to FLAG IP to enrich FLAG-K-Ras4a-G12V or -G12V-3KR with its interacting proteins. After washing the FLAG resin five times with IP washing buffer, the resins from ‘heavy’ and ‘light’ cells were mixed. Enriched proteins on the resin were eluted with triple FLAG peptide following the manufacturer’s protocol. Eluted proteins were precipitated with methanol/chloroform/water (4/1.5/3 volume ratio with the sample volume set as 1), and the protein pellets were washed twice with 1 mL ice-cold methanol. The protein pellets were air dried for 10-15 min, and subjected to disulfide reduction and protein denaturation in 100 μL of buffer containing 6 M urea, 10 mM DTT and 50 mM Tris-HCl pH 8.0 at room temperature for 1 h. Then iodoacetamide (final concentration 40 mM) was added to alkylate the proteins at room temperature for 1 h. Subsequently, DTT (final concentration 40 mM) was added to stop alkylation at room temperature for 1 h. The samples were then diluted 7 times with buffer containing 1 mM CaCl_2_ and 50 mM Tris-HCl pH 8.0 and digested with 2 μg trypsin at 37°C for 12 h. Trypsin digestion was quenched with 0.2 % trifluoroacetic acid. Then the mixture was desalted using Sep-Pak C18 cartridge following the manufacturer's protocol and subjected to liquid chromatography (LC)-MS/MS analysis.

The lyophilized peptides were reconstituted in 2% acetonitrile (ACN) with 0.5% formic acid (FA) and analyzed by LTQ-Orbitrap Elite mass spectrometer coupled with nanoLC. Reconstituted peptides were injected onto Acclaim PepMap nano Viper C18 trap column (5 μm, 100 μm × 2 cm, Thermo Dionex) for online desalting and then separated on C18 RP nano column (5 μm, 75 μm × 50 cm, Magic C18, Bruker). The flow rate was 0.3 μL/min, and the gradient was 5-38% ACN with 0.1% FA from 0120 min, 38-95% ACN with 0.1% FA from 120-127 min, and 95% ACN with 0.1% FA from 127-135 min. The Orbitrap Elite was operated in positive ion mode with spray voltage 1.6 kV and source temperature 275 °C. Data-dependent acquisition (DDA) mode was used by one precursor ions MS survey scan from m/z 375 to 1800 at resolution 120,000 using FT mass analyzer, followed by up to 10 MS/MS scans at resolution 15,000 on 10 most intensive peaks. Collision-induced dissociation (CID) parameters were set with isolation width 2.0 m/z and normalized collision energy at 35%. All data were acquired in Xcalibur 2.2 operation software. MS1 and MS2 data were processed using Sequest HT software within the Proteome Discoverer 1.4.1.14 (PD 1.4, Thermo Scientific).

### Detection of lysine fatty acylation on Ras by mass spectrometry (MS)

To detect H-Ras lysine fatty acylation, HEK293T cells were transfected with pCMV5-H-Ras for 24 h and treated with 50 μM Alk14 for another 6 h. To detect K-Ras4a lysine fatty acylation, HEK293T cells with stable SIRT2 KD were transfected with pCMV5-FLAG-K-Ras4a for 24 h and treated with or without 50 μM Alk14 for another 6 h. Cells were collected and lysed in 1% NP-40 lysis buffer with protease inhibitor cocktail. FLAG IP was then performed with 50 mg of total protein lysate to purify FLAG-K-Ras4a or Flag-H-Ras. After washing the FLAG resin three times with IP washing buffer, H-Ras or K-Ras4a was eluted by heating at 95 °C for 10 min in buffer containing 1% SDS and 50 mM Tris-HCl pH 8.0. After centrifuging at 15,000 g for 2 min, the supernatant was transferred to a new tube and was treated with 300 mM NH_2_OH pH 7.4 at 95 °C for 10 min. The Ras protein was then precipitated by methanol/chloroform and processed (disulfide reduction, denaturing, alkylation and neutralization) as described above. The resultant Ras protein was digested with 2 μg of trypsin at 37°C for 2 h in a glass vial (to avoid absorption of the fatty acylated peptide by plastics). Then desalting was done using Sep-Pak C18 cartridge following the manufacturer’s protocol.

For the LC-MS/MS analysis of the digested peptides, the same settings described for the SILAC experiment were applied except the LC gradient was 5-95% ACN with 0.1% FA from 0-140 min. The settings for identifying Alk14 modification in Sequest were: two miscleavages for full trypsin with fixed carbamidomethyl modification of cysteine residue, dynamic modifications of 234.198 Da (Alk14) on lysine residue, N-terminal acetylation, methionine oxidation and deamidation of asparagine and glutamine residues. The peptide mass tolerance and fragment mass tolerance values were 15 p.p.m. and 0. 8 Da, respectively.

### Detection of lysine fatty acylation on endogenous Ras in HCT116 cells

HCT116 cells (parental cells, or cells infected with shCtrl/shSIRT2-carrying lentivirus for 3 days) were cultured with fresh medium containing 50 μM Alk14 for 6 h. Cells were collected and lysed using the same method described above. Pan-Ras immunoprecipitation was performed using pan-Ras (Y13-259) antibody by following manufacturer's protocol. The lysine fatty acylation on endogenous Ras was detected by on-beads click chemistry and in-gel fluorescence using the same method described above. To directly detect lysine fatty acylation on endogenous Ras by MS, 200 mg of total lysates from HCT116 cells with SIRT2 KD was used for pan-RAS immunoprecipitation, followed by denaturation, alkylation, neutralization, trypsin digestion and LC-MS/MS analysis using the same method described above.

### Statistical analysis

Quantitative imaging data were expressed in box plot as indicated in figure legends. Statistical evaluation of imaging data was done using two-way ANOVA. Other quantitative data were expressed in scatter plots with mean ± SEM (standard error of the mean, shown as error bar) shown. Differences between two groups were examined using unpaired two-tailed Student’s t test. The *P* values were indicated (\**P* < 0.05, ** *P* < 0.01, and *** *P* < 0.001). *P* values < 0.05 were considered statistically significant. No statistical tool was used to pre-determine sample size. No blinding was done, no randomization was used, and no sample was excluded from analysis.

## Supplemental Information

**Figure S1.**
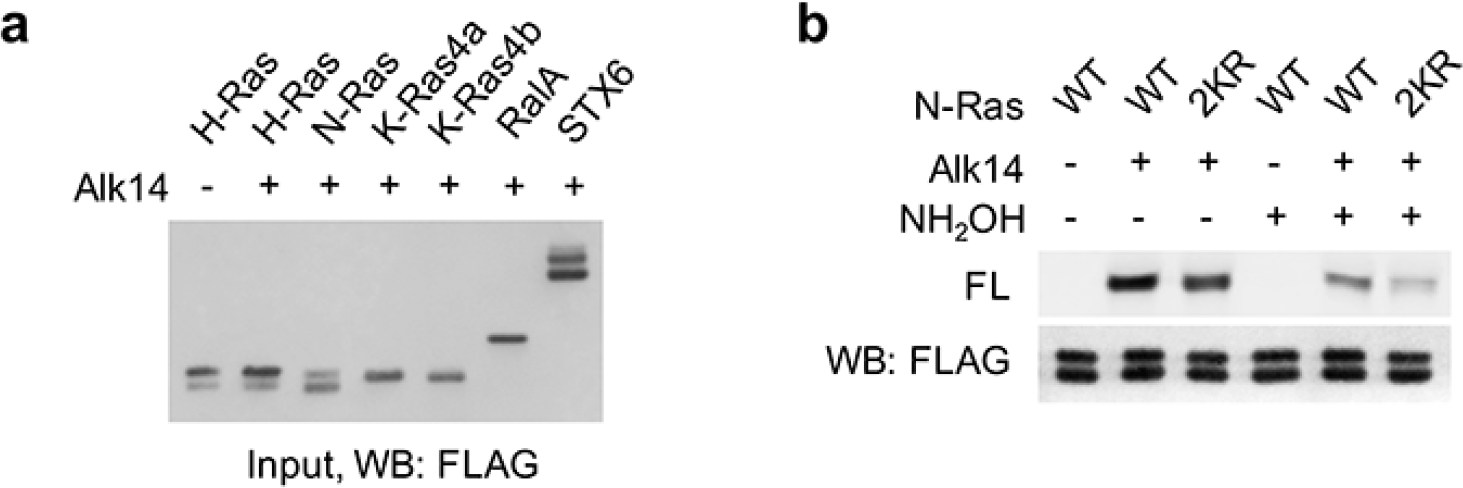
N-Ras proteins may be lysine fatty acylated. (**a**) Representative western blot analyses of FLAG-tagged Ras protein, RalA and STX6 in whole cell extracts. (**b**) In-gel fluorescence showing the fatty acylation levels of N-Ras WT and 2KR mutant without or with NH2OH treatment. Representative images from three independent experiments are shown.

**Figure S2.**
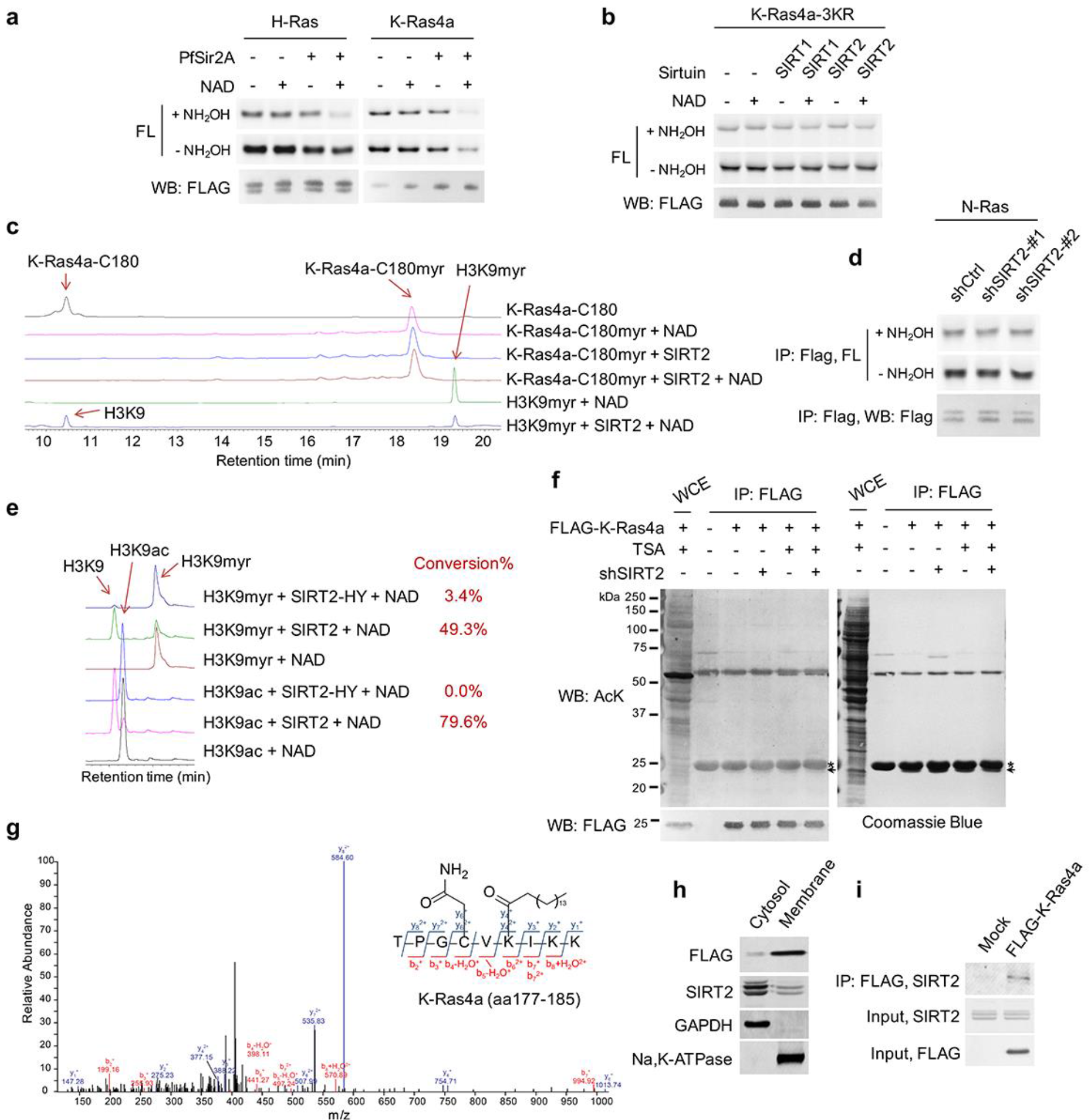
SIRT2 regulates lysine fatty acylation of K-Ras4a. (**a**) In-gel fluorescence detection of fatty acylation on H-Ras and K-Ras4a treated without or with 10μM PfSir2A and 1 mM NAD *in vitro.* (**b**) Fatty acylation of K-Ras4a-3KR treated without or with 5 μM SIRT2 and 1 mM NAD *in vitro.* (**c**) High-performance liquid chromatography (HPLC) traces showing SIRT2 hydrolyzing myristoyl group from H3K9myr peptide but not K-Ras4a-C180myr peptide. The reaction with H3K9myr, SIRT2 and NAD serves as a control to show that SIRT2 was active. (**d**) Effect of SIRT2 KD on the fatty acylation level of N-Ras in HEK293T cells (**e**) Comparison of the activities of SIRT2-WT and SIRT2-HY on H3K9ac and H3K9myr peptides by HPLC-based *in vitro* assay. The conversion rates are shown on the right. (**f**) Acetylation of K-Ras4a in Ctrl or SIRT2 KD (by shSIRT2-#2) HEK293T cells treated with ethanol or TSA (1 μM) for 1 h. The “*” points to the light chain of the anti-FLAG antibody, while the arrow points to K-Ras4a. (**g**) MS/MS spectrum of triply charged K-Ras4a peptide with palmitoylation on K182. The b- and y-ions are shown along with the peptide sequence. The cysteine residue was carbamidomethylated due to iodoacetamide treatment during sample preparation. (**h**) Subcellular fractionation showing the localization of SIRT2 and FLAG-K-Ras4a. (**i**) Co-IP of FLAG-K-Ras4a with endogenous SIRT2 in HEK293T cells. Representative images from three independent experiments are shown.

**Figure S3.**
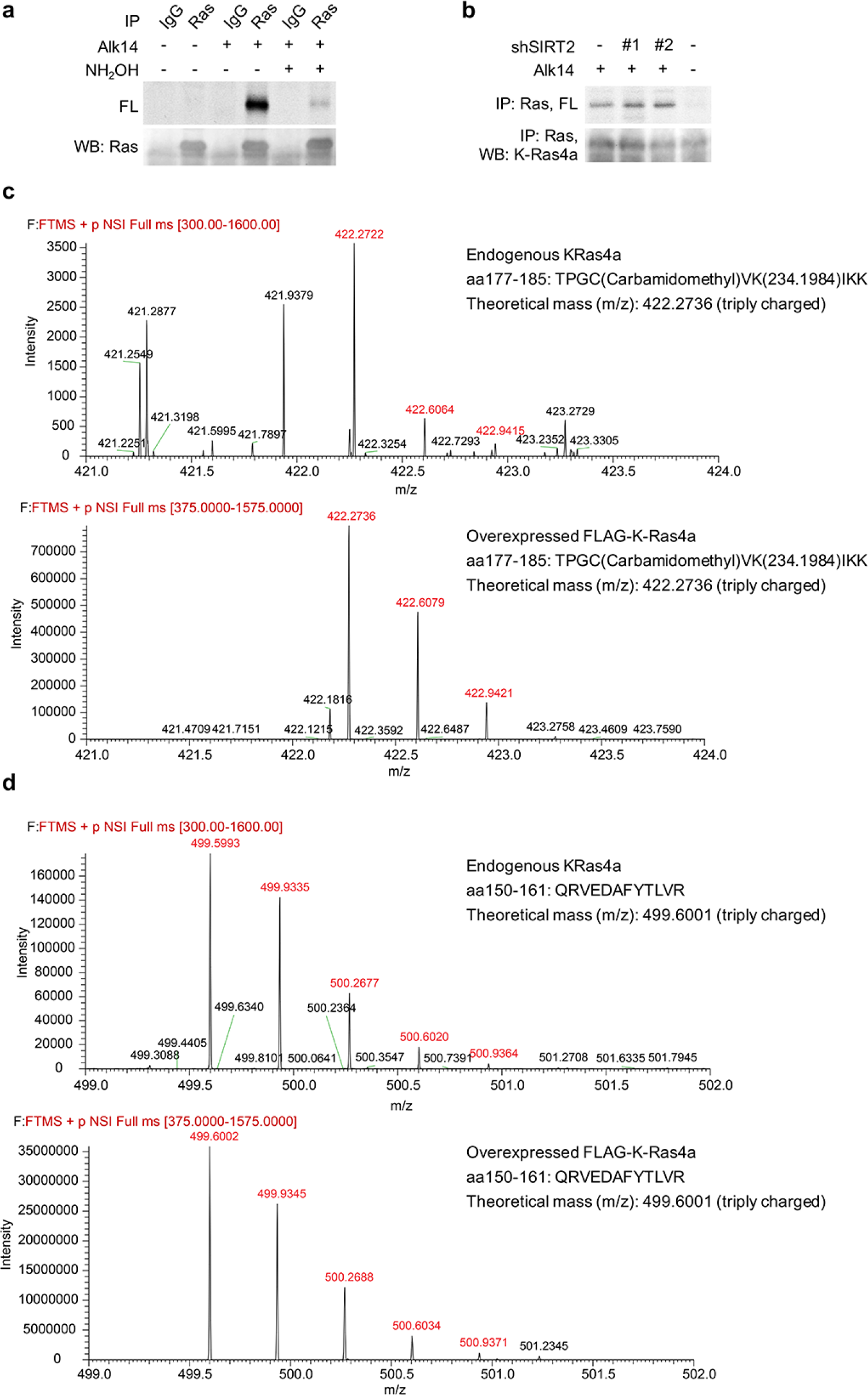
Endogenous K-Ras4a is lysine fatty acylated. (**a**) In-gel fluorescence detection of fatty acylation on endogenous total Ras proteins immunoprecipiated from HCT116 cells. (**b**) Effect of SIRT2 KD on fatty acylation of endogenous Ras after NH_2_OH treatment. (**c**) Comparison of the MS spectra of Alk14-modified K-Ras4a aa177-185 peptide from endogenous Ras and overexpressed K-Ras4a MS analyses. (**d**) Comparison of the MS spectra of K-Ras4a aa 150-161 peptide from endogenous Ras and overexpressed K-Ras4a MS analyses. The ion intensities for the Alk14-modified aa177-185 and the unmodified aa 150-161 peptides were over 200 times lower than those from overexpressed K-Ras4a.

**Figure S4.**
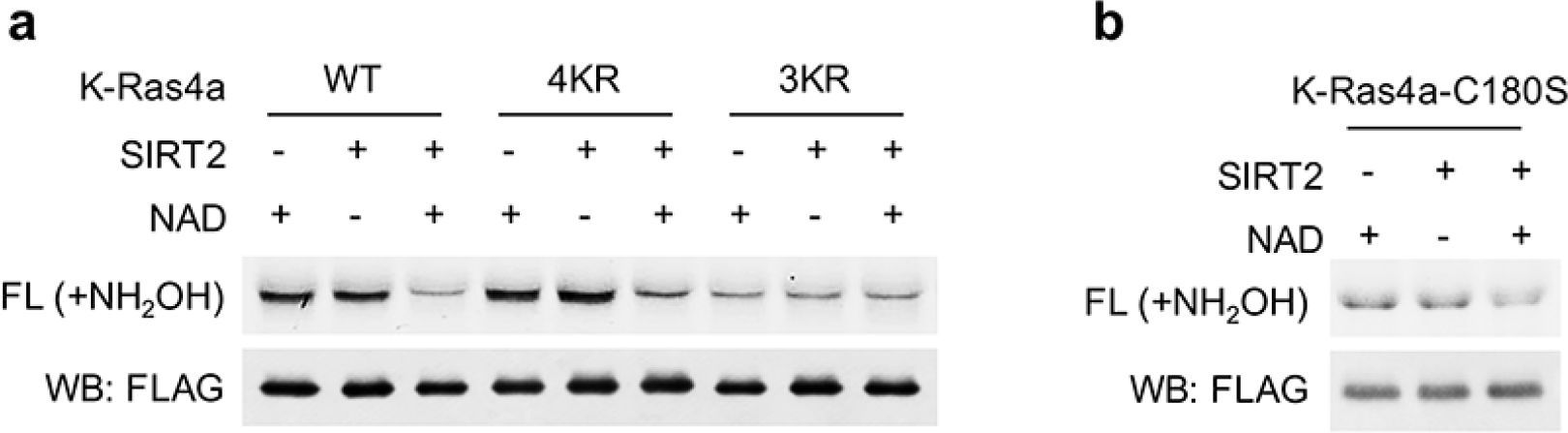
SIRT2 removes lysine fatty acylation from K-Ras4a-WT, -4KR and -C180S, but not -3KR. Representative images showing in-gel fluorescence detection of fatty acylation of K-Ras4a-WT, -4KR, -3KR (a) and -C180S (b) treated without or with 5 μM SIRT2 and 1 mM NAD *in vitro.* Representative images from three independent experiments are shown.

**Figure S5.**
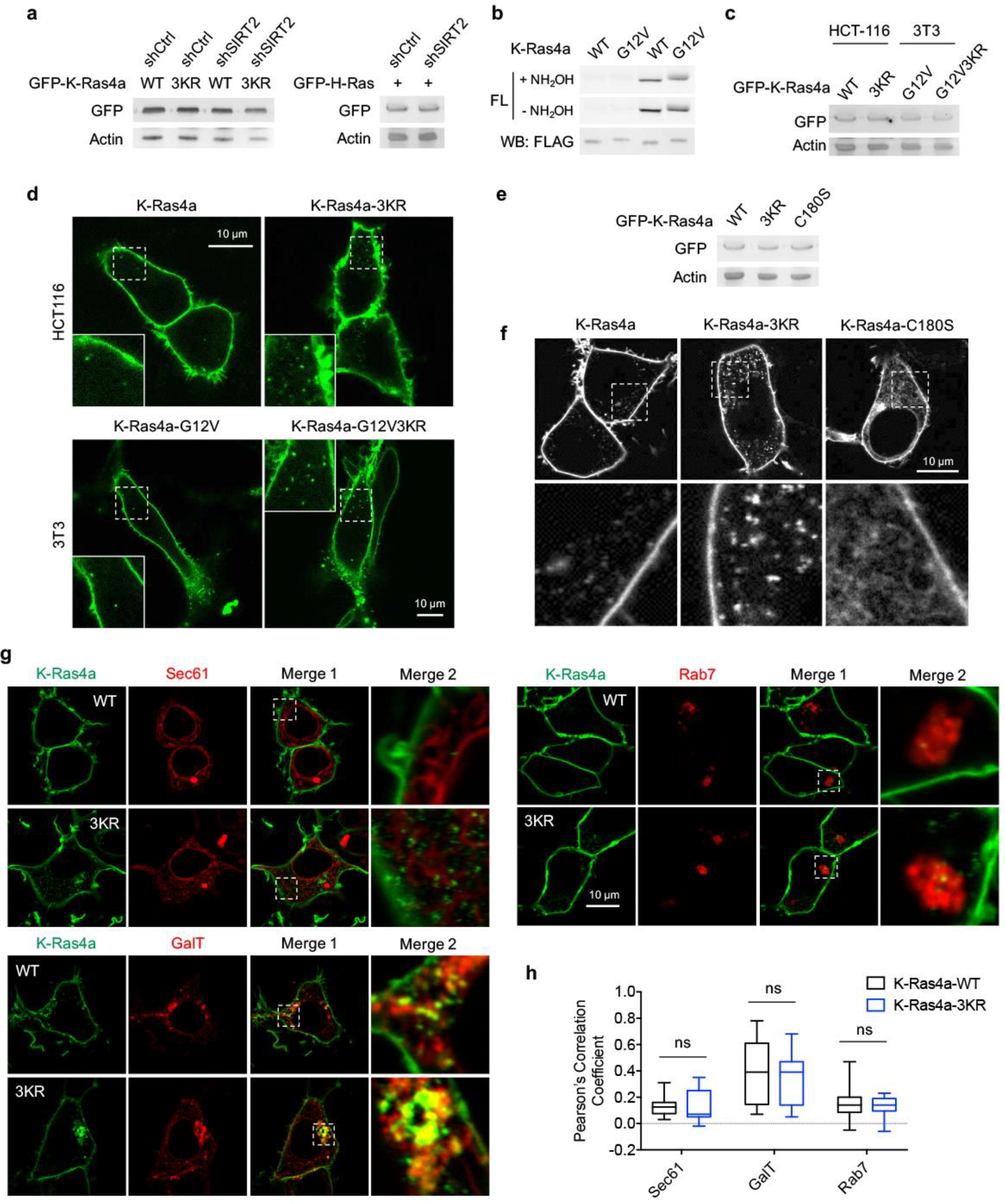
Lysine fatty acylation regulates subcellular localization of K-Ras4a. (**a**) Western blot analyses showing equal overexpression of GFP-K-Ras4a WT and 3KR in HEK293T cells with Ctrl and SIRT2 KD. (**b**) Fatty acylation levels of K-Ras4a WT and G12V in HEK293T cells. (**c**) Western blot analyses showing equal protein levels for overexpressed GFP-K-Ras4a-WT and --3KR in HCT116 cells, and GFP-K-Ras4a-G12V and -G12V-3KR in 3T3 cells. (**d**) Live cell imaging of HCT116 cells overexpressing GFP-K-Ras4a-WT and -3KR and NIH 3T3 cells overexpressing GFP-K-Ras4a-G12V and -G12V-3KR. Insets are magnifications of the regions enclosed by the white dashed squares. (**e**) Western blot showing equal protein levels for overexpressed GFP-K-Ras4a-WT, -3KR and -C180S in HEK293T cells. (**f**) Confocal images showing subcellular localization of GFP-K-Ras4a, -3KR, and -C180S. The bottom panels show the magnified images of the regions enclosed by the white dashed squares in the top panels. (**g**) Representative images for examining the colocalization of GFP-K-Ras4a or -3KR with Sec61, GalT, and Rab7 in HEK293T cells. Magnifications of the white dashed squares-enclosed regions in Merge 1 are shown as Merge 2. (**h**) Statistical analyses of the colocalization from (g) using Pearson's coefficient (n= 11, 11, 11, 11, 13, 13 cells for each sample from left to right, respectively). The images shown are representative of 80100% of the cells examined. Statistical evaluation was done using two-way ANOVA. Centre line of the box plot represents the mean value, box represents the 95 % confidence interval, and whiskers represent the range of the values. ns, not significant.

**Figure S6.**
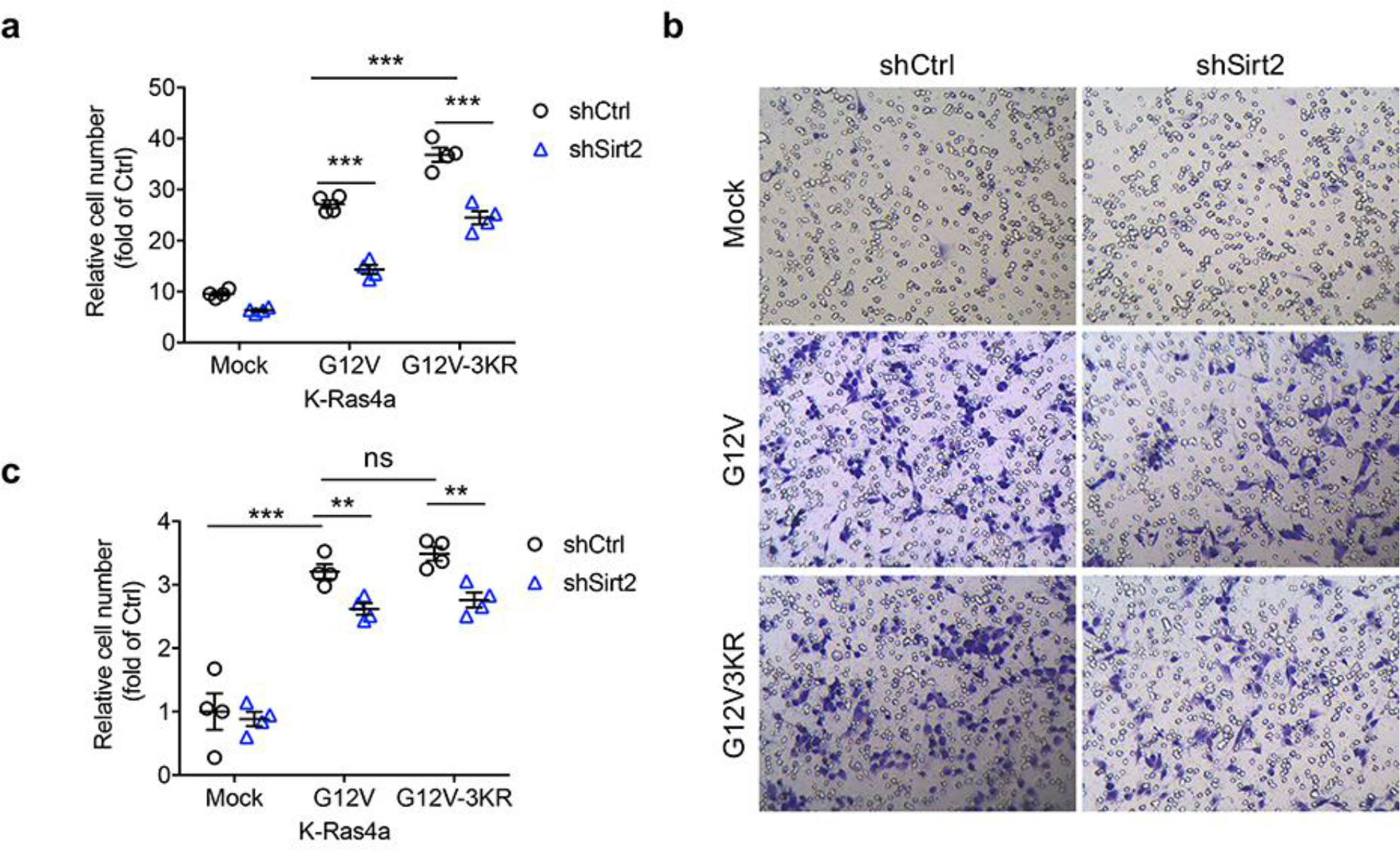
Lysine fatty acylation regulates K-Ras4a-G12V-mediated cell proliferation but not migration. (**a**) Effect of Ctrl or Sirt2 KD on proliferation of NIH3T3 cells stably overexpressing Mock, K-Ras4a-G12V or -G12V-3KR. Cell numbers were determined by crystal violet staining 0 or 5 days after the transduction with shCtrl or shSirt2-carrying lentivirus. The y axis represents cell numbers normalized to that of the corresponding shCtrl group on Day 0. (**b**) Representative images of transwell migration assay in NIH3T3 cells stably overexpressing Mock, K-Ras4a-G12V or -G12V-3KR with Ctrl or Sirt2 KD. (**c**) Migration cell numbers in (**b**) relative to that of Mock with Ctrl KD. Statistical evaluation was done using unpaired two-tailed Student’s t test. Error bars represent SEM in four biological replicates. \*\**P* < 0.01; \*\*\**P* < 0.001; ns, not significant.

**Figure S7.**
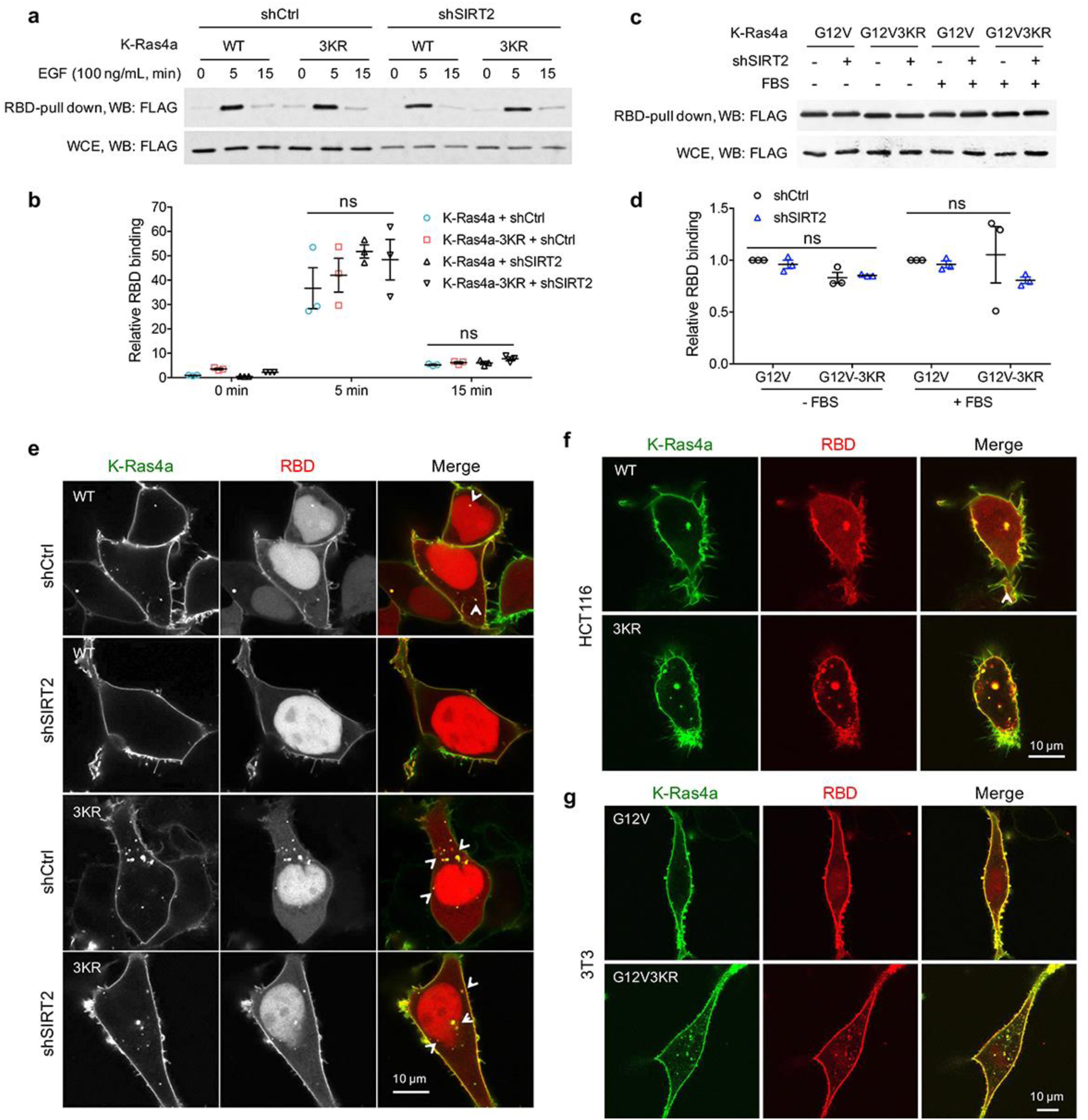
Lysine fatty acylation regulates the subcellular localization of active K-Ras4a. (**a**, **b**) RBD pull-down assay in HEK293T cells expressing FLAG-K-Ras4a WT or 3KR with Ctrl or SIRT2 KD (by shSIRT2-#2) (a). Cells were serum-starved overnight and treated with 100 ng/mL EGF for 0, 5 and 15 min. The relative RBD binding with respect to cells expressing K-Ras4a and shCtrl at 0 min was quantified in (b). (**c**, **d**) RBD pull-down assay in HEK293T cells expressing FLAG-K-Ras4a-G12V or -G12V-3KR with Ctrl or SIRT2 KD (c). Cells were cultured in FBS-free or complete medium for 12 h before being subjected to RBD pull-down. RBD binding relative to cells expressing K-Ras4a-G12V-shCtrl was quantified in (d). (**e**) Co-localization of GFP-K-Ras4a WT or 3KR with DsRed-RBD in live HEK293T cells with Ctrl or SIRT2 KD. (**f**) Live cell imaging showing the colocalization of GFP-K-Ras4a WT or 3KR with DsRed-RBD in HCT116 cells. (**g**) Live cell imaging showing the colocalization of GFP-K-Ras4a-G12V or -G12V-3KR with DsRed-RBD in NIH3T3 cells. Error bars represent SEM in three biological replicates. The images shown are representative of 80-100% of the cells examined. Statistical evaluation was done using unpaired two-tailed Student’s t test. Error bars represent SEM in four biological replicates. ns, not significant.

**Figure S8.**
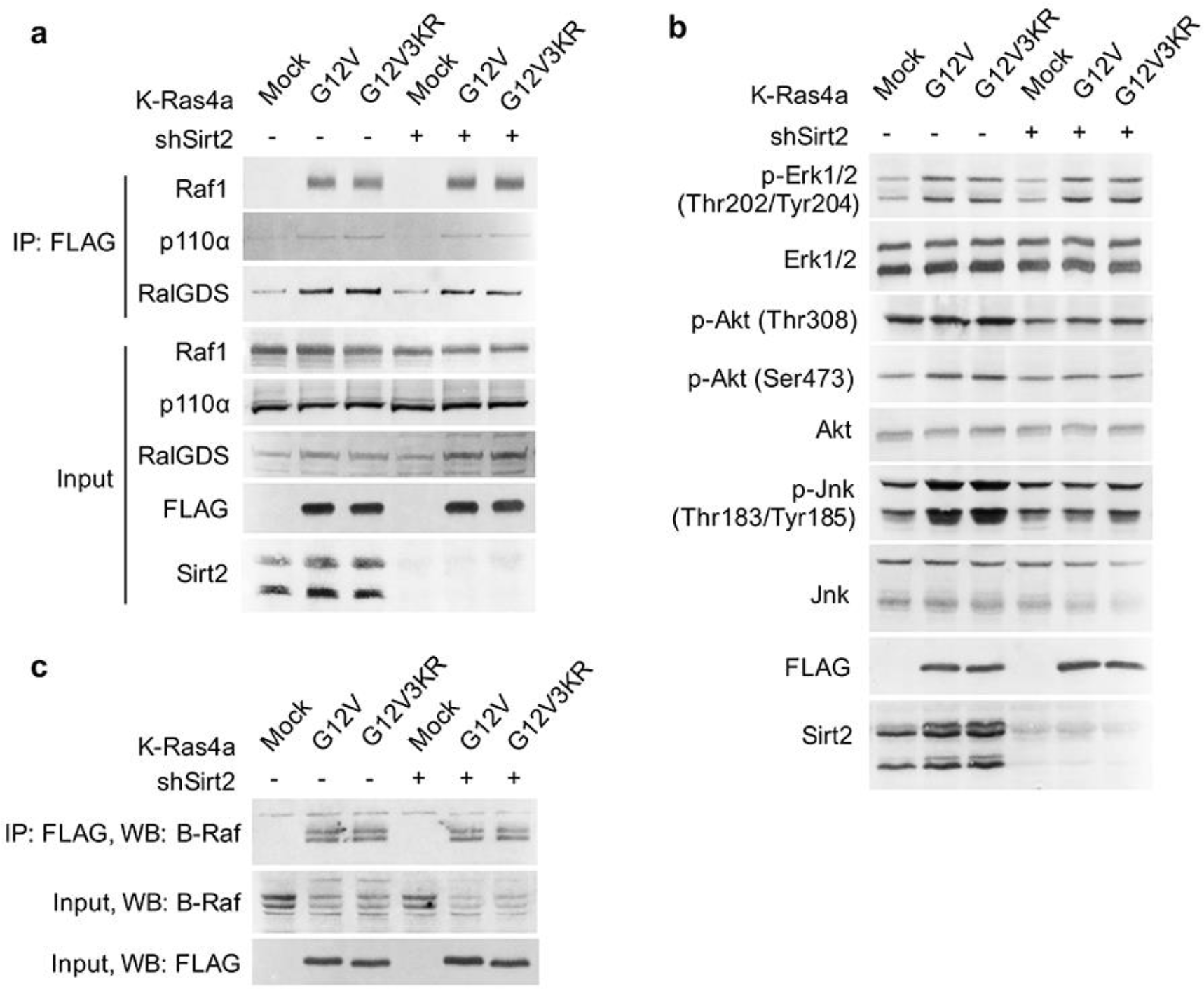
Lysine fatty acylation does not affect K-Ras4a signaling through Raf1, PI3K, RalGDS or B-Raf. (**a**) Co-IP of FLAG and Raf1, p110a or RalGDS in NIH3T3 cells stably expressing Mock, FLAG-K-Ras4a-G12V or -G12V-3KR with Ctrl or Sirt2 KD. (**b**) Western blot analyses of phospho-Erk, -Akt and -Jnk in NIH3T3 cells stably expressing Mock, FLAG-K-Ras4a-G12V or -G12V-3KR with Ctrl or Sirt2 KD. (**c**) Co-IP of FLAG and B-Raf in NIH3T3 cells stably expressing Mock, FLAG-K-Ras4a-G12V or FLAG-K-Ras4a-G12V-3KR with Ctrl or Sirt2 KD. Representative images from three independent experiments are shown.

**Figure S9.**
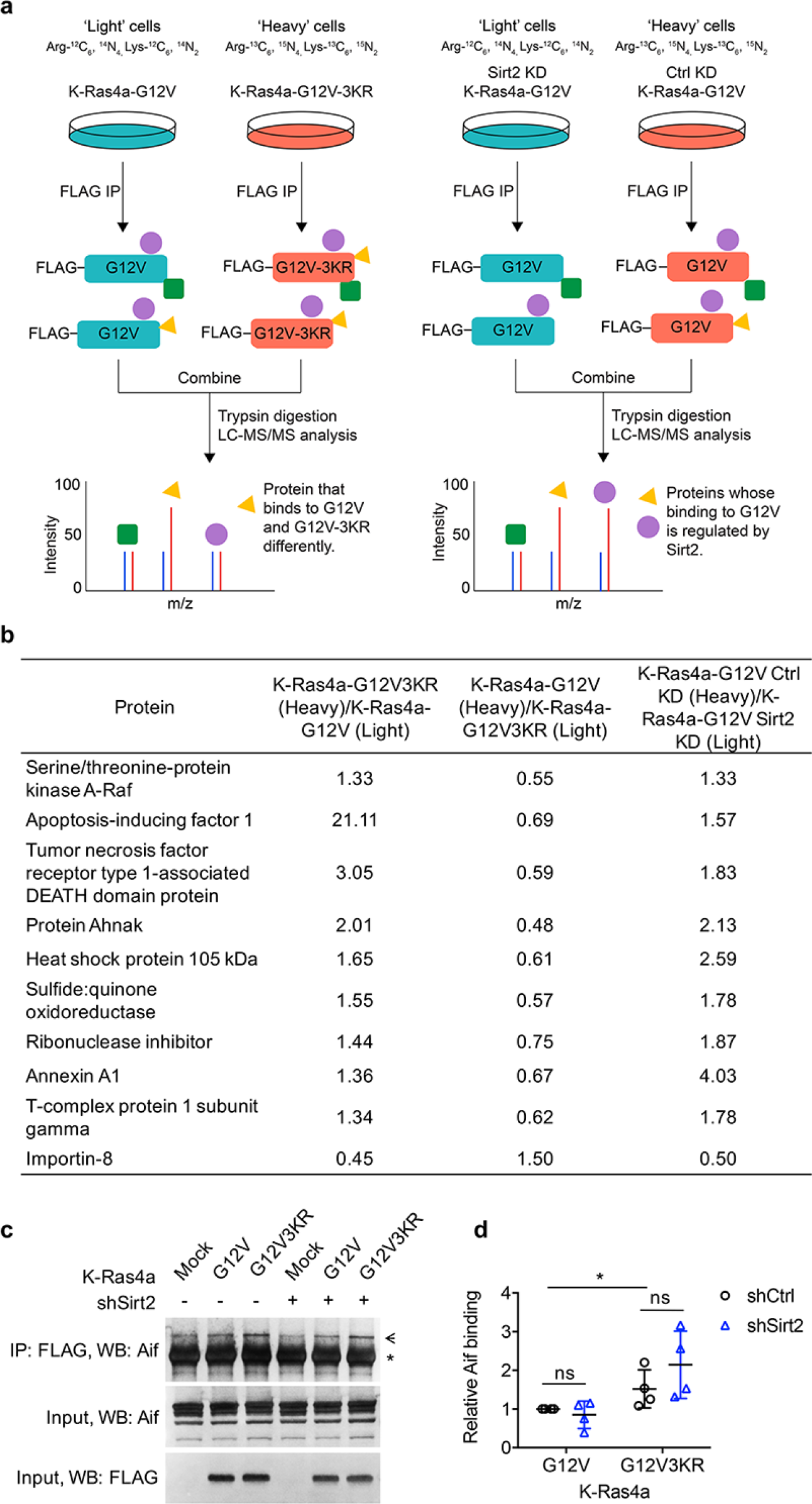
Interactome study identifies K-Ras4a-G12V interacting proteins that potentially mediate the effect of lysine fatty acylation. (**a**) Schematic workflow of the K-Ras4a-G12V SILAC interactome study. (**b**) List of proteins whose binding to K-Ras4a-G12V may be altered (H/L > 1.3 or < 0.77) by lysine fatty acylation. (**c**) Co-IP of FlAg and Aif in NIH3T3 cells stably expressing Mock, FLAG-K-Ras4a-G12V or -G12V-3KR with Ctrl or Sirt2 KD. The “*” points to the heavy chain of the anti-FLAG antibody, while the arrow points to Aif. (**d**) Quantification of relative Aif binding level in (**c**) compared to that in cells expressing K-Ras4a-G12V-shCtrl. Statistical evaluation was by unpaired two-tailed Student’s t test. Error bars represent SEM in four biological replicates. \**P* < 0.05; ns, not significant. Representative images from four independent experiments are shown.

**Figure S10.**
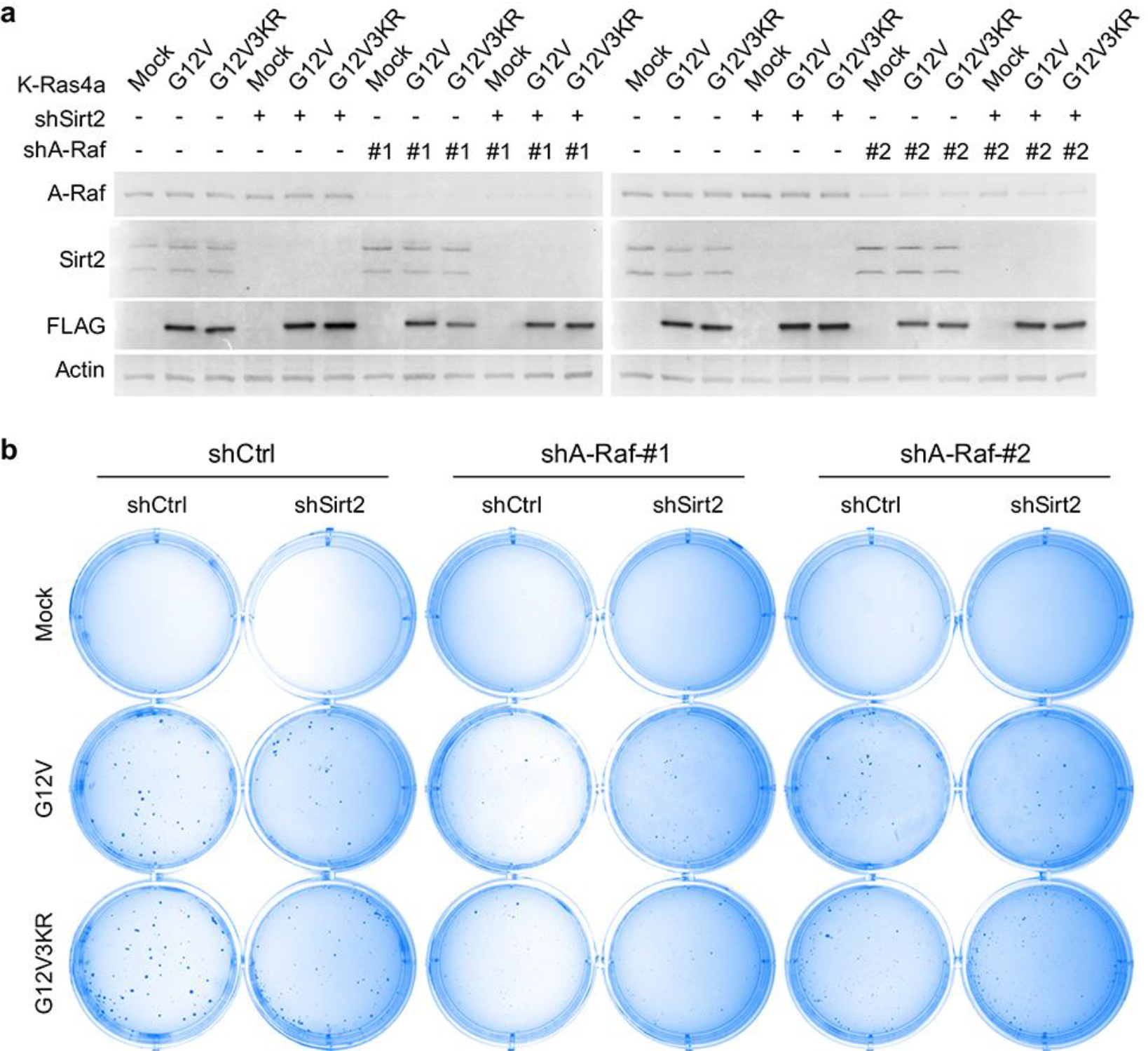
A-Raf mediates the regulation of K-Ras4a-G12V transforming activity by lysine fatty acylation. (**a**) Western blot analysis of A-Raf, Sirt2 and FLAG in NIH3T3 cells expressing Mock, FLAG-K-Ras4a-G12V or -G12V-3KR with Ctrl or SIRT2 KD, and Ctrl or A-Raf KD. (**b**) Images showing anchorage-independent growth of NIH3T3 cells stably expressing the K-Ras4a-G12V or -G12V-3KR with Ctrl or Sirt2 KD, and Ctrl or A-Raf KDs. Representative images from three independent experiments are shown.

**Figure S11.**
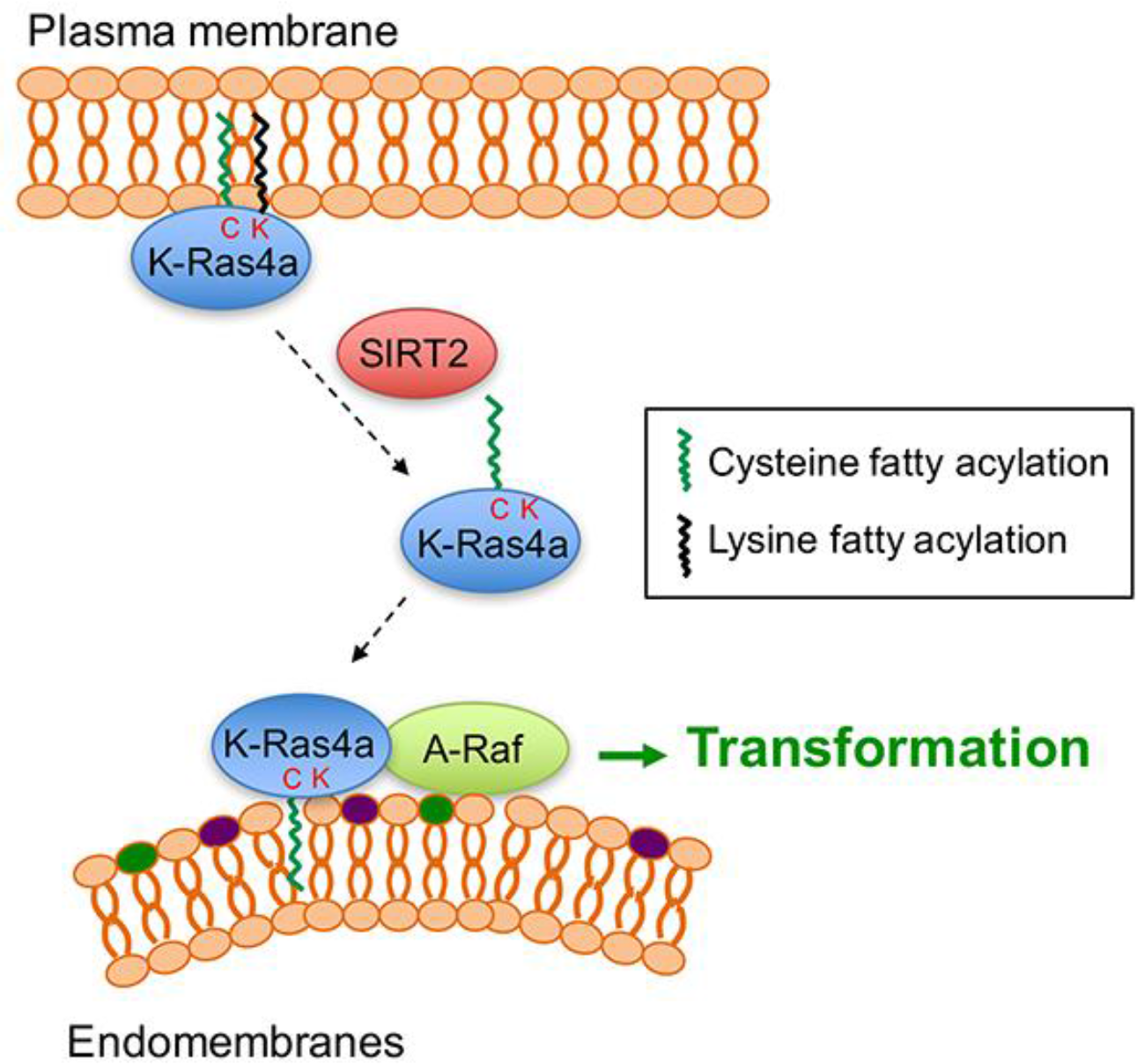
Model for the regulation of K-Ras4a by SIRT2-mediated removal of lysine fatty acylation. Removal of lysine fatty acylation by SIRT2 facilitates K-Ras4a to localize to endomembranes and interact with A-Raf, and thus enhances its activity to transform cells.

**Figure S12.**
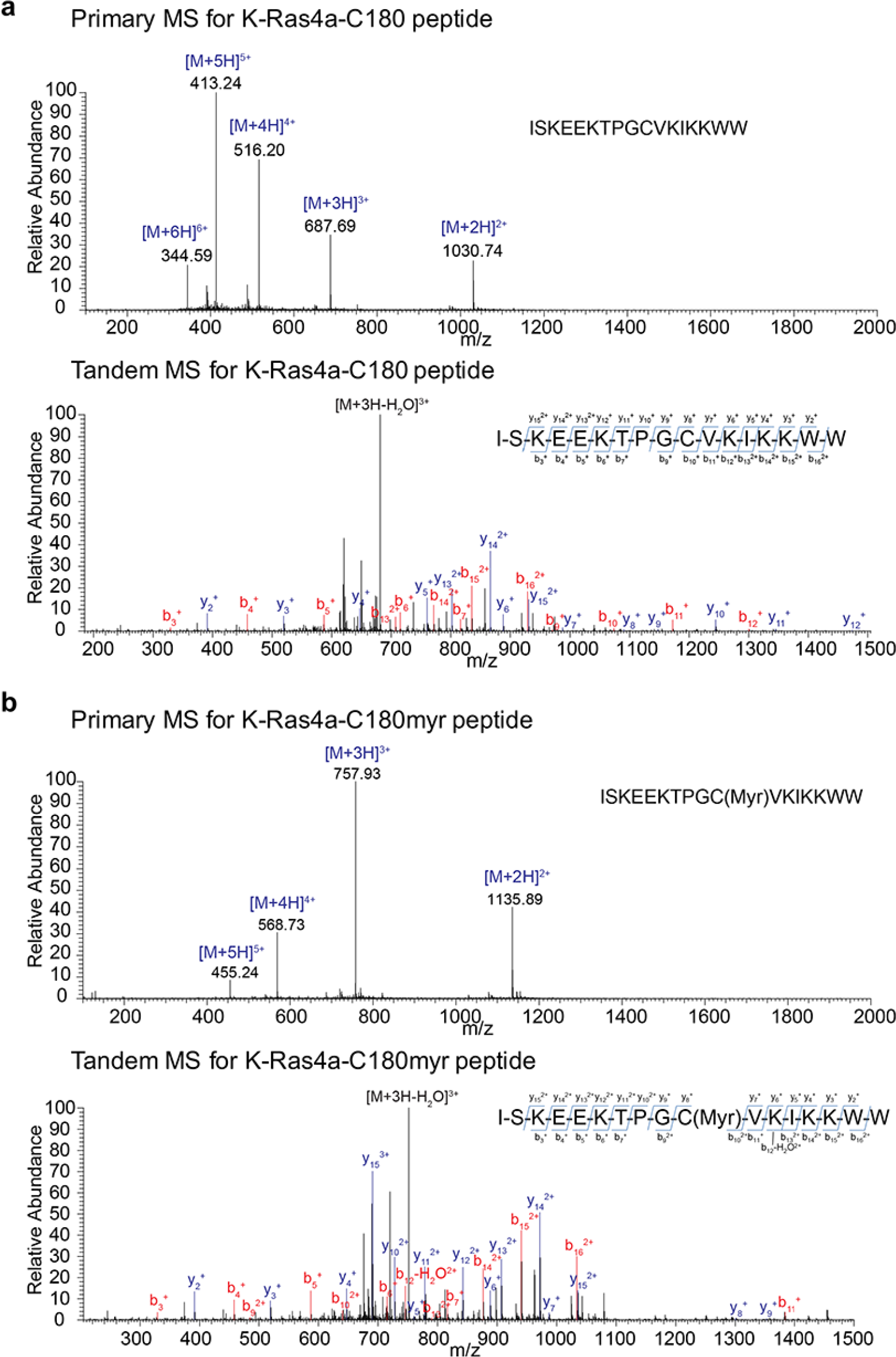
MS and tandem MS/MS spectra for the synthetic free K-Ras4a-C180 (a) and myristoyl K-Ras4a-C180 (b) peptides.

**Movie S1.** Dynamics of K-Ras4a in HEK293T cells. Cells were transfected with GFP-K-Ras4a for 24 h before being subjected to time-lapse confocal microscopy. Images were collected at 1-s intervals for 1 min of a single plane.

**Movie S2.** Dynamics of K-Ras4a-3KR in HEK293T cells. Cells were transfected with GFP-K-Ras4a-3KR for 24 h before being subjected to time-lapse confocal microscopy. Images were collected at 1-s intervals for 1 min of a single plane.

**Movie S3.** Dynamics of K-Ras4a-C180S in HEK293T cells. Cells were transfected with GFP-K-Ras4a-3KR for 24 h before being subjected to time-lapse confocal microscopy. Images were collected at 1-s intervals for 1 min of a single plane.

## References

1. Peng, T., Thinon, E. & Hang, H.C. Proteomic analysis of fatty-acylated proteins. Curr Opin Chem Biol 30, 77–86 (2016).

2. Tate, E.W., Kalesh, K.A., Lanyon-Hogg, T., Storck, E.M. & Thinon, E. Global profiling of protein lipidation using chemical proteomic technologies. Curr Opin Chem Biol 24, 48–57 (2015).

3. Lanyon-Hogg, T., Faronato, M., Serwa, R.A. & Tate, E.W. Dynamic Protein Acylation: New Substrates, Mechanisms, and Drug Targets. Trends Biochem Sci 42, 566–581 (2017).

4. Hedo, J.A., Collier, E. & Watkinson, A. Myristyl and palmityl acylation of the insulin receptor. J Biol Chem 262, 954–7 (1987).

5. Stevenson, F.T., Bursten, S.L., Fanton, C., Locksley, R.M. & Lovett, D.H. The 31-kDa precursor of interleukin 1 alpha is myristoylated on specific lysines within the 16-kDa N-terminal propiece. Proc Natl Acad Sci U S A 90, 7245–9 (1993).

6. Bursten, S.L., Locksley, R.M., Ryan, J.L. & Lovett, D.H. Acylation of monocyte and glomerular mesangial cell proteins. Myristyl acylation of the interleukin 1 precursors. J Clin Invest 82, 1479–88 (1988).

7. Pillai, S. & Baltimore, D. Myristoylation and the post-translational acquisition of hydrophobicity by the membrane immunoglobulin heavy-chain polypeptide in B lymphocytes. Proc Natl Acad Sci U S A 84, 7654–8 (1987).

8. Stevenson, F.T., Bursten, S.L., Locksley, R.M. & Lovett, D.H. Myristyl acylation of the tumor necrosis factor alpha precursor on specific lysine residues. J Exp Med 176, 1053–62 (1992).

9. Jiang, H., Khan, S., Wang, Y., Charron, G., He, B., Sebastian, C., Du, J., Kim, R., Ge, E., Mostoslavsky, R., Hang, H.C., Hao, Q. & Lin, H. SIRT6 regulates TNF-a secretion through hydrolysis of long-chain fatty acyl lysine. Nature 496, 110–113 (2013).

10. Jiang, H., Zhang, X. & Lin, H. Lysine fatty acylation promotes lysosomal targeting of TNF-alpha. Sci Rep 6, 24371 (2016).

11. Zhang, X., Khan, S., Jiang, H., Antonyak, M.A., Chen, X., Spiegelman, N.A., Shrimp, J.H., Cerione, R.A. & Lin, H. Identifying the functional contribution of the defatty-acylase activity of SIRT6. Nat Chem Biol 12, 614–20 (2016).

12. Feldman, J.L., Baeza, J. & Denu, J.M. Activation of the protein deacetylase SIRT6 by long-chain fatty acids and widespread deacylation by mammalian sirtuins. J Biol Chem 288, 31350–6 (2013).

13. Bao, X., Wang, Y., Li, X., Li, X.M., Liu, Z., Yang, T., Wong, C.F., Zhang, J., Hao, Q. & Li, X.D. Identification of ‘erasers' for lysine crotonylated histone marks using a chemical proteomics approach. Elife 3(2014).

14. Teng, Y.B., Jing, H., Aramsangtienchai, P., He, B., Khan, S., Hu, J., Lin, H. & Hao, Q. Efficient demyristoylase activity of SIRT2 revealed by kinetic and structural studies. Sci Rep 5, 8529 (2015).

15. Liu, Z., Yang, T., Li, X., Peng, T., Hang, H.C. & Li, X.D. Integrative chemical biology approaches for identification and characterization of “erasers” for fatty-acid-acylated lysine residues within proteins. Angew Chem Int Ed Engl 54, 1149–52 (2015).

16. He, B., Hu, J., Zhang, X. & Lin, H. Thiomyristoyl peptides as cell-permeable Sirt6 inhibitors. Org. Biomol. Chem. 12, 7498–502 (2014).

17. Malumbres, M. & Barbacid, M. RAS oncogenes: the first 30 years. Nat Rev Cancer 3, 459–65 (2003).

18. Hancock, J.F. Ras proteins: different signals from different locations. Nat Rev Mol Cell Biol 4, 373–84 (2003).

19. Tsai, F.D., Lopes, M.S., Zhou, M., Court, H., Ponce, O., Fiordalisi, J.J., Gierut, J.J., Cox, A.D., Haigis, K.M. & Philips, M.R. K-Ras4A splice variant is widely expressed in cancer and uses a hybrid membrane-targeting motif. Proc Natl Acad Sci U S A (2015).

20. Zhao, H., Liu, P., Zhang, R., Wu, M., Li, D., Zhao, X., Zhang, C., Jiao, B., Chen, B., Chen, Z. & Ren, R. Roles of palmitoylation and the KIKK membrane-targeting motif in leukemogenesis by oncogenic KRAS4A. J Hematol Oncol 8, 132 (2015).

21. To, M.D., Wong, C.E., Karnezis, A.N., Del Rosario, R., Di Lauro, R. & Balmain, A. Kras regulatory elements and exon 4A determine mutation specificity in lung cancer. Nat Genet 40, 1240–4 (2008).

22. Nishimura, A. & Linder, M.E. Identification of a novel prenyl and palmitoyl modification at the CaaX motif of Cdc42 that regulates RhoGDI binding. Mol Cell Biol 33, 1417–29 (2013).

23. Kang, R., Wan, J., Arstikaitis, P., Takahashi, H., Huang, K., Bailey, A.O., Thompson, J.X., Roth, A.F., Drisdel, R.C., Mastro, R., Green, W.N., Yates, J.R., 3rd, Davis, N.G. & El-Husseini, A. Neural palmitoyl-proteomics reveals dynamic synaptic palmitoylation. Nature 456, 904–9 (2008).

24. Swaney, D.L., Wenger, C.D. & Coon, J.J. Value of using multiple proteases for large-scale mass spectrometry-based proteomics. J Proteome Res 9, 1323–9 (2010).

25. Zhang, X., Spiegelman, N.A., Nelson, O.D., Jing, H. & Lin, H. SIRT6 regulates Ras-related protein R-Ras2 by lysine defatty-acylation. Elife 6(2017).

26. Zhu, A.Y., Zhou, Y., Khan, S., Deitsch, K.W., Hao, Q. & Lin, H. Plasmodium falciparum Sir2A preferentially hydrolyzes medium and long chain fatty acyl lysine. ACS Chem Biol 7, 155–9 (2012).

27. North, B.J. & Verdin, E. Mitotic regulation of SIRT2 by cyclin-dependent kinase 1-dependent phosphorylation. J Biol Chem 282, 19546–55 (2007).

28. Hoff, K.G., Avalos, J.L., Sens, K. & Wolberger, C. Insights into the sirtuin mechanism from ternary complexes containing NAD+ and acetylated peptide. Structure 14, 1231–40 (2006).

29. Jackson, M.D., Schmidt, M.T., Oppenheimer, N.J. & Denu, J.M. Mechanism of nicotinamide inhibition and transglycosidation by Sir2 histone/protein deacetylases. J Biol Chem 278, 50985–98 (2003).

30. Du, J., Zhou, Y., Su, X., Yu, J., Khan, S., Jiang, H., Kim, J., Woo, J., Kim, J.H., Choi, B.H., He, B., Chen, W., Zhang, S., Cerione, R.A., Auwerx, J., Hao, Q. & Lin, H. Sirt5 is an NAD-dependent protein lysine demalonylase and desuccinylase. Science 334, 806–809 (2011).

31. Wilson, J.P., Raghavan, A.S., Yang, Y.-Y., Charron, G. & Hang, H.C. Proteomic analysis of fatty-acylated proteins in mammalian cells with chemical reporters reveals S-acylation of histone H3 variants. Mol. Cell. Proteomics 10(2011).

32. Omary, M.B. & Trowbridge, I.S. Covalent binding of fatty acid to the transferrin receptor in cultured human cells. J Biol Chem 256, 4715–8 (1981).

33. Lampson, B.L., Pershing, N.L., Prinz, J.A., Lacsina, J.R., Marzluff, W.F., Nicchitta, C.V., MacAlpine, D.M. & Counter, C.M. Rare codons regulate KRas oncogenesis. Curr Biol 23, 70–5 (2013).

34. Ali, M., Kaltenbrun, E., Anderson, G.R., Stephens, S.J., Arena, S., Bardelli, A., Counter, C.M. & Wood, K.C. Codon bias imposes a targetable limitation on KRAS-driven therapeutic resistance. Nat Commun 8, 15617 (2017).

35. North, B.J. & Verdin, E. Interphase nucleo-cytoplasmic shuttling and localization of SIRT2 during mitosis. PLoS One 2, e784 (2007).

36. Inoue, T., Hiratsuka, M., Osaki, M., Yamada, H., Kishimoto, I., Yamaguchi, S., Nakano, S., Katoh, M., Ito, H. & Oshimura, M. SIRT2, a tubulin deacetylase, acts to block the entry to chromosome condensation in response to mitotic stress. Oncogene 26, 945–57 (2007).

37. Wright, L.P. & Philips, M.R. Thematic review series: lipid posttranslational modifications. CAAX modification and membrane targeting of Ras. J Lipid Res 47, 883–91 (2006).

38. Choy, E., Chiu, V.K., Silletti, J., Feoktistov, M., Morimoto, T., Michaelson, D., Ivanov, I.E. & Philips, M.R. Endomembrane trafficking of ras: the CAAX motif targets proteins to the ER and Golgi. Cell 98, 69–80 (1999).

39. Rocks, O., Peyker, A., Kahms, M., Verveer, P.J., Koerner, C., Lumbierres, M., Kuhlmann, J., Waldmann, H., Wittinghofer, A. & Bastiaens, P.I. An acylation cycle regulates localization and activity of palmitoylated Ras isoforms. Science 307, 1746–52 (2005).

40. Ballester, R., Furth, M.E. & Rosen, O.M. Phorbol ester-and protein kinase C-mediated phosphorylation of the cellular Kirsten ras gene product. J Biol Chem 262, 2688–95 (1987).

41. Bivona, T.G., Quatela, S.E., Bodemann, B.O., Ahearn, I.M., Soskis, M.J., Mor, A., Miura, J., Wiener, H.H., Wright, L., Saba, S.G., Yim, D., Fein, A., Perez de Castro, I., Li, C., Thompson, C.B., Cox, A.D. & Philips, M.R. PKC regulates a farnesyl-electrostatic switch on K-Ras that promotes its association with Bcl-XL on mitochondria and induces apoptosis. Mol Cell 21, 481–93 (2006).

42. Jura, N., Scotto-Lavino, E., Sobczyk, A. & Bar-Sagi, D. Differential modification of Ras proteins by ubiquitination. Mol Cell 21, 679–87 (2006).

43. Chiu, V.K., Bivona, T., Hach, A., Sajous, J.B., Silletti, J., Wiener, H., Johnson, R.L., 2nd, Cox, A.D. & Philips, M.R. Ras signalling on the endoplasmic reticulum and the Golgi. Nat Cell Biol 4, 343–50 (2002).

44. Lu, A., Tebar, F., Alvarez-Moya, B., Lopez-Alcala, C., Calvo, M., Enrich, C., Agell, N., Nakamura, T., Matsuda, M. & Bachs, O. A clathrin-dependent pathway leads to KRas signaling on late endosomes en route to lysosomes. J Cell Biol 184, 863–79 (2009).

45. Howe, C.L., Valletta, J.S., Rusnak, A.S. & Mobley, W.C. NGF signaling from clathrin-coated vesicles: evidence that signaling endosomes serve as a platform for the Ras-MAPK pathway. Neuron 32, 801–14 (2001).

46. Misaki, R., Morimatsu, M., Uemura, T., Waguri, S., Miyoshi, E., Taniguchi, N., Matsuda, M. & Taguchi, T. Palmitoylated Ras proteins traffic through recycling endosomes to the plasma membrane during exocytosis. J Cell Biol 191, 23–9 (2010).

47. Apolloni, A., Prior, I.A., Lindsay, M., Parton, R.G. & Hancock, J.F. H-ras but not K-ras traffics to the plasma membrane through the exocytic pathway. Mol Cell Biol 20, 2475–87 (2000).

48. Fivaz, M. & Meyer, T. Reversible intracellular translocation of KRas but not HRas in hippocampal neurons regulated by Ca2+/calmodulin. J Cell Biol 170, 429–41 (2005).

49. Grant, B.D. & Donaldson, J.G. Pathways and mechanisms of endocytic recycling. Nat Rev Mol Cell Biol 10, 597–608 (2009).

50. Zhou, W., Ni, T.K., Wronski, A., Glass, B., Skibinski, A., Beck, A. & Kuperwasser, C. The SIRT2 Deacetylase Stabilizes Slug to Control Malignancy of Basal-like Breast Cancer. Cell Rep 17, 1302–1317 (2016).

51. He, X., Nie, H., Hong, Y., Sheng, C., Xia, W. & Ying, W. SIRT2 activity is required for the survival of C6 glioma cells. Biochem. Biophys. Res. Comm. 417, 468–472 (2012).

52. Liu, P.Y., Xu, N., Malyukova, A., Scarlett, C.J., Sun, Y.T., Zhang, X.D., Ling, D., Su, S.P., Nelson, C., Chang, D.K., Koach, J., Tee, A.E., Haber, M., Norris, M.D., Toon, C., Rooman, I., Xue, C., Cheung, B.B., Kumar, S., Marshall, G.M., Biankin, A.V. & Liu, T. The histone deacetylase SIRT2 stabilizes Myc oncoproteins. Cell Death Differ. 20, 503–514 (2013).

53. Jing, H., Hu, J., He, B., Negron Abril, Y.L., Stupinski, J., Weiser, K., Carbonaro, M., Chiang, Y.L., Southard, T., Giannakakou, P., Weiss, R.S. & Lin, H. A SIRT2-Selective Inhibitor Promotes c-Myc Oncoprotein Degradation and Exhibits Broad Anticancer Activity. Cancer Cell 29, 297–310 (2016).

54. Zhao, D., Zou, S.-W., Liu, Y., Zhou, X., Mo, Y., Wang, P., Xu, Y.-H., Dong, B., Xiong, Y., Lei, Q.-Y. & Guan, K.-L. Lysine-5 acetylation negatively regulates lactate dehydrogenase A and is decreased in pancreatic cancer. Cancer cell 23, 464–476 (2013).

55. Jing, H. & Lin, H. Sirtuins in epigenetic regulation. Chem Rev 115, 2350–75 (2015).

56. Hu, J., Jing, H. & Lin, H. Sirtuin inhibitors as anticancer agents. Future Med Chem 6, 945–66 (2014).

57. Wang, Y.P., Zhou, L.S., Zhao, Y.Z., Wang, S.W., Chen, L.L., Liu, L.X., Ling, Z.Q., Hu, F.J., Sun, Y.P., Zhang, J.Y., Yang, C., Yang, Y., Xiong, Y., Guan, K.L. & Ye, D. Regulation of G6PD acetylation by SIRT2 and KAT9 modulates NADPH homeostasis and cell survival during oxidative stress. EMBO J 33, 1304–20 (2014).

58. Xu, S.N., Wang, T.S., Li, X. & Wang, Y.P. SIRT2 activates G6PD to enhance NADPH production and promote leukaemia cell proliferation. Sci Rep 6, 32734 (2016).

59. Dekker, F.J., Rocks, O., Vartak, N., Menninger, S., Hedberg, C., Balamurugan, R., Wetzel, S., Renner, S., Gerauer, M., Scholermann, B., Rusch, M., Kramer, J.W., Rauh, D., Coates, G.W., Brunsveld, L., Bastiaens, P.I. & Waldmann, H. Small-molecule inhibition of APT1 affects Ras localization and signaling. Nat Chem Biol 6, 449–56 (2010).

60. Chandra, A., Grecco, H.E., Pisupati, V., Perera, D., Cassidy, L., Skoulidis, F., Ismail, S.A., Hedberg, C., Hanzal-Bayer, M., Venkitaraman, A.R., Wittinghofer, A. & Bastiaens, P.I. The GDI-like solubilizing factor PDEdelta sustains the spatial organization and signalling of Ras family proteins. Nat Cell Biol 14, 148–58 (2011).

61. Berndt, N., Hamilton, A.D. & Sebti, S.M. Targeting protein prenylation for cancer therapy. Nat Rev Cancer 11, 775–91 (2011).

62. Rauch, J., O'Neill, E., Mack, B., Matthias, C., Munz, M., Kolch, W. & Gires, O. Heterogeneous nuclear ribonucleoprotein H blocks MST2-mediated apoptosis in cancer cells by regulating A-Raf transcription. Cancer Res 70, 1679–88 (2010).

63. Rauch, J., Vandamme, D., Mack, B., McCann, B., Volinsky, N., Blanco, A., Gires, O. & Kolch, W. Differential localization of A-Raf regulates MST2-mediated apoptosis during epithelial differentiation. Cell Death Differ 23, 1283–95 (2016).

64. Lee, W., Jiang, Z., Liu, J., Haverty, P.M., Guan, Y., Stinson, J., Yue, P., Zhang, Y., Pant, K.P., Bhatt, D., Ha, C., Johnson, S., Kennemer, M.I., Mohan, S., Nazarenko, I., Watanabe, C., Sparks, A.B., Shames, D.S., Gentleman, R., de Sauvage, F.J., Stern, H., Pandita, A., Ballinger, D.G., Drmanac, R., Modrusan, Z., Seshagiri, S. & Zhang, Z. The mutation spectrum revealed by paired genome sequences from a lung cancer patient. Nature 465, 473–7 (2010).

65. Imielinski, M., Greulich, H., Kaplan, B., Araujo, L., Amann, J., Horn, L., Schiller, J., Villalona-Calero, M.A., Meyerson, M. & Carbone, D.P. Oncogenic and sorafenib-sensitive ARAF mutations in lung adenocarcinoma. J Clin Invest 124, 1582–6 (2014).

66. Nelson, D.S., Quispel, W., Badalian-Very, G., van Halteren, A.G., van den Bos, C., Bovee, J.V., Tian, S.Y., Van Hummelen, P., Ducar, M., MacConaill, L.E., Egeler, R.M. & Rollins, B.J. Somatic activating ARAF mutations in Langerhans cell histiocytosis. Blood 123, 3152–5 (2014).

67. Hangen, E., Blomgren, K., Benit, P., Kroemer, G. & Modjtahedi, N. Life with or without AIF. Trends Biochem Sci 35, 278–87 (2010).

68. van den Gupta, Y., Grabocka, E. & Bar-Sagi, D. RAS oncogenes: weaving a tumorigenic web. Nat Rev Cancer 11, 761–74 (2011).

69. Roy, S., Wyse, B. & Hancock, J.F. H-Ras signaling and K-Ras signaling are differentially dependent on endocytosis. Mol Cell Biol 22, 5128–40 (2002).

70. An, S., Yang, Y., Ward, R., Liu, Y., Guo, X.X. & Xu, T.R. A-Raf: A new star of the family of raf kinases. Crit Rev Biochem Mol Biol 50, 520–31 (2015).

71. Buss, J.E. & Sefton, B.M. Direct identification of palmitic acid as the lipid attached to p21ras. Mol Cell Biol 6, 116–22 (1986).

72. Hancock, J.F., Magee, A.I., Childs, J.E. & Marshall, C.J. All ras proteins are polyisoprenylated but only some are palmitoylated. Cell 57, 1167–77 (1989).

73. Drisdel, R.C. & Green, W.N. Labeling and quantifying sites of protein palmitoylation. Biotechniques 36, 276–85 (2004).

74. Forrester, M.T., Hess, D.T., Thompson, J.W., Hultman, R., Moseley, M.A., Stamler, J.S. & Casey, P.J. Site-specific analysis of protein S-acylation by resin-assisted capture. J Lipid Res 52, 393–8 (2011).

75. Zhou, Y., Prakash, P., Liang, H., Cho, K.J., Gorfe, A.A. & Hancock, J.F. Lipid-Sorting Specificity Encoded in K-Ras Membrane Anchor Regulates Signal Output. Cell 168, 239–251 e16 (2017).

76. Goodwin, J.S., Drake, K.R., Rogers, C., Wright, L., Lippincott-Schwartz, J., Philips, M.R. & Kenworthy, A.K. Depalmitoylated Ras traffics to and from the Golgi complex via a nonvesicular pathway. J Cell Biol 170, 261–72 (2005).

77. Wellbrock, C., Karasarides, M. & Marais, R. The RAF proteins take centre stage. Nat Rev Mol Cell Biol 5, 875–85 (2004).

78. Matallanas, D., Birtwistle, M., Romano, D., Zebisch, A., Rauch, J., von Kriegsheim, A. & Kolch, W. Raf family kinases: old dogs have learned new tricks. Genes Cancer 2, 232–60 (2011).

79. Weber, C.K., Slupsky, J.R., Herrmann, C., Schuler, M., Rapp, U.R. & Block, C. Mitogenic signaling of Ras is regulated by differential interaction with Raf isozymes. Oncogene 19, 169–76 (2000).

80. Williams, J.G., Drugan, J.K., Yi, G.S., Clark, G.J., Der, C.J. & Campbell, S.L. Elucidation of binding determinants and functional consequences of Ras/Raf-cysteine-rich domain interactions. J Biol Chem 275, 22172–9 (2000).

81. Fischer, A., Hekman, M., Kuhlmann, J., Rubio, I., Wiese, S. & Rapp, U.R. B- and C-RAF display essential differences in their binding to Ras: the isotype-specific N terminus of B-RAF facilitates Ras binding. J Biol Chem 282, 26503–16 (2007).

82. Cox, A.D., Fesik, S.W., Kimmelman, A.C., Luo, J. & Der, C.J. Drugging the undruggable RAS: Mission possible? Nat Rev Drug Discov 13, 828–51 (2014).

83. Shimizu, K., Birnbaum, D., Ruley, M.A., Fasano, O., Suard, Y., Edlund, L., Taparowsky, E., Goldfarb, M. & Wigler, M. Structure of the Ki-ras gene of the human lung carcinoma cell line Calu-1. Nature 304, 497–500 (1983).

84. Bheda, P., Jing, H., Wolberger, C. & Lin, H. The Substrate Specificity of Sirtuins. Annu Rev Biochem (2016).

85. Wilking-Busch, M.J., Ndiaye, M.A., Huang, W. & Ahmad, N. Expression Profile of SIRT2 in Human Melanoma and Implications for Sirtuin-Based Chemotherapy. Cell Cycle 0 (2017).

86. Shah, A.A., Ito, A., Nakata, A. & Yoshida, M. Identification of a Selective SIRT2 Inhibitor and Its Antibreast Cancer Activity. Biol Pharm Bull 39, 1739–1742 (2016).

87. Moniot, S., Forgione, M., Lucidi, A., Hailu, G.S., Nebbioso, A., Carafa, V., Baratta, F., Altucci, L., Giacche, N., Passeri, D., Pellicciari, R., Mai, A., Steegborn, C. & Rotili, D. Development of 1,2,4-Oxadiazoles as Potent and Selective Inhibitors of the Human Deacetylase Sirtuin 2: Structure-Activity Relationship, X-Ray Crystal Structure and Anticancer Activity. J Med Chem (2017).

88. Charron, G., Zhang, M.M., Yount, J.S., Wilson, J., Raghavan, A.S., Shamir, E. & Hang, H.C. Robust fluorescent detection of protein fatty-acylation with chemical reporters. J Am Chem Soc 131, 4967–75 (2009).

89. Liu, H. & Naismith, J.H. An efficient one-step site-directed deletion, insertion, single and multiple-site plasmid mutagenesis protocol. BMC Biotechnol 8, 91 (2008).

90. Luo, X., Wasilko, D.J., Liu, Y., Sun, J., Wu, X., Luo, Z.Q. & Mao, Y. Structure of the Legionella Virulence Factor, SidC Reveals a Unique PI(4) P-Specific Binding Domain Essential for Its Targeting to the Bacterial Phagosome. PLoS Pathog 11, e1004965 (2015).

91. Zurek, N., Sparks, L. & Voeltz, G. Reticulon short hairpin transmembrane domains are used to shape ER tubules. Traffic 12, 28–41 (2011).

92. Choudhury, A., Dominguez, M., Puri, V., Sharma, D.K., Narita, K., Wheatley, C.L., Marks, D.L. & Pagano, R.E. Rab proteins mediate Golgi transport of caveola-internalized glycosphingolipids and correct lipid trafficking in Niemann-Pick C cells. J Clin Invest 109, 1541–50 (2002).

93. Sherer, N.M., Lehmann, M.J., Jimenez-Soto, L.F., Ingmundson, A., Horner, S.M., Cicchetti, G., Allen, P.G., Pypaert, M., Cunningham, J.M. & Mothes, W. Visualization of retroviral replication in living cells reveals budding into multivesicular bodies. Traffic 4, 785–801 (2003).

94. Schindelin, J., Arganda-Carreras, I., Frise, E., Kaynig, V., Longair, M., Pietzsch, T., Preibisch, S., Rueden, C., Saalfeld, S., Schmid, B., Tinevez, J.Y., White, D.J., Hartenstein, V., Eliceiri, K., Tomancak, P. & Cardona, A. Fiji: an open-source platform for biological-image analysis. Nat Methods 9, 676–82 (2012).

95. Du, J., Jiang, H. & Lin, H. Investigating the ADP-ribosyltransferase activity of sirtuins with NAD analogues and 32P-NAD. Biochemistry 48, 2878–90 (2009).

96. Adler, J. & Parmryd, I. Quantifying colocalization by correlation: the Pearson correlation coefficient is superior to the Mander's overlap coefficient. Cytometry A 77, 733–42 (2010).

